# Endothelin receptor blockade potentiates adoptive T cell and CAR T cell therapies

**DOI:** 10.64898/2026.09.02.748239

**Authors:** Charys Papagregoriou, Fotios Mpekris, Jongwon Yoon, Georgios Kastrappis, Andria A. Lytridou, Maria Thrasyvoulou, Konstantinos Symeonidis, Christina Pitsilidou, Constantina Neophytou, Myrofora Panagi, Chrysovalantis Voutouri, Maria Iakovou, Gregoria Gregoriou, Triantafyllos Stylianopoulos, Paul Maciocia, Paul Costeas, Philippos Demetriou

## Abstract

Adoptive T cell therapies have transformed the treatment of selected haematological malignancies but remain limited in many cancer settings by biological barriers that restrict effective and durable antitumour responses. Here, we investigated whether pharmacological blockade of the endothelin receptor pathway, which regulates vascular, stromal and immune processes relevant to antitumour responses, could enhance T cell-based cellular immunotherapy. Using orthotopic 4T1 triple-negative breast cancer and A20 B cell lymphoma models, we evaluated endothelin receptor blockade with non-engineered adoptive T cell therapy and anti-CD19 CAR T cell therapy, respectively, with or without immune checkpoint inhibition. Endothelin receptor blockade substantially increased tumour control responses across both therapeutic platforms, while the addition of immune checkpoint inhibition further increased their frequency and promoted maintained complete responses, resulting in the most durable tumour control. Mice achieving maintained complete responses resisted tumour rechallenge, consistent with long-term antitumour immune protection. High-dimensional T cell profiling identified distinct intratumoral T cell states associated with tumour control and elevated CD2 expression as a recurring feature across these populations. Together, these findings identify endothelin receptor blockade as a rational combinatorial strategy for enhancing T cell-based cellular immunotherapy and provide a strong rationale for the clinical evaluation of this therapeutic strategy.

## INTRODUCTION

T cell-based cellular immunotherapies have transformed cancer treatment, with the capacity to induce durable clinical responses and long-term remissions, particularly in haematological malignancies. This therapeutic approach encompasses multiple platforms, including non-engineered adoptive T cell therapies, such as tumour-infiltrating lymphocytes (TILs) and tumour-draining lymph node (TDLN)-derived T cells, as well as genetically engineered T cells expressing tumour-specific T cell receptors (TCRs) or chimeric antigen receptors (CARs)^1–6^. Non-engineered adoptive T cell therapy (ACT), particularly TIL therapy, provided some of the earliest clinical evidence that adoptively transferred tumour-reactive T cells can mediate profound and durable tumour regression in patients with advanced melanoma^7,8^. The clinical efficacy of TIL therapy in advanced melanoma has since been further demonstrated in a randomised phase III trial^9^. Among engineered approaches, CAR T cell therapy has demonstrated remarkable clinical efficacy in several B-cell malignancies, largely due to the accessibility of malignant B cells and the availability of well-defined target antigens^10–12^. These advances have established engineered T cell therapies as a major therapeutic modality for haematological cancers. More recently, clinical studies have demonstrated that CAR T cell therapy can also mediate clinically meaningful antitumour activity in selected solid tumour settings, including mesothelin-targeted CAR T cells in malignant pleural disease and GD2-directed CAR T cells in diffuse midline glioma^13,14^.

Despite these advances and encouraging clinical activity in selected indications, durable and consistent responses to T cell-based cellular immunotherapy remain uncommon across most solid tumours^15,16^. Therapeutic success depends on overcoming multiple complementary biological and physical barriers that collectively influence the induction, magnitude and durability of effective antitumour T cell responses. These include tumour-intrinsic mechanisms^17–19^, stromal^20,21^ and vascular barriers^22–25^, immunosuppressive features of the tumour microenvironment ranging from soluble mediators such as TGF-β^26,27^ to immune checkpoint pathways such as PD-1/PD-L1 and CTLA-4^28–30^, together with factors that influence the functional fitness and persistence of the transferred T cell product^31–36^. Collectively, these barriers limit the therapeutic efficacy of T cell-based cellular immunotherapy across many solid tumour settings.

Among the therapeutic strategies developed to overcome these barriers, immune checkpoint inhibitors (ICIs), such as those targeting PD-1/PD-L1 and CTLA-4, have been combined with both non-engineered and engineered adoptive T cell therapies and shown to enhance transferred T cell function, persistence and antitumour activity^37–40^. Despite these improvements, both the magnitude and durability of therapeutic benefit remain variable, with many tumours retaining resistance despite the addition of ICI. This suggests that maximising therapeutic efficacy will likely require targeting complementary biological processes beyond inhibitory checkpoint signalling alone.

Among the pathways that may offer complementary therapeutic opportunities, the endothelin (ET) axis has emerged as a potential target. The endothelin system comprises a family of vasoactive peptides and their cognate receptors, endothelin receptor A (ETA) and endothelin receptor B (ETB). Beyond its established roles in tumour cell proliferation, survival, invasion and metastasis^41^, endothelin signalling regulates multiple aspects of tumour biology, including vascular function, immune cell trafficking and function^23,42,43^, stromal architecture and fibrosis^42,44^, as well as inflammatory responses^45^. We therefore hypothesised that pharmacological inhibition of the endothelin receptor pathway could enhance the efficacy of T cell-based cellular immunotherapy by alleviating biological barriers limiting effective antitumour T cell responses. Supporting the biological rationale for this hypothesis, ETB signalling in the tumour endothelium has been shown to establish a barrier to T cell homing in a preclinical tumour vaccine model^23^, while pharmacological blockade of both ETA and ETB has been shown to improve tumour perfusion, alter tumour mechanical properties and increase T cell representation within tumours^42^. Further studies have identified ETB signalling as a regulator of the pro-angiogenic and immunosuppressive functions of a distinct iron-rich tumour-associated macrophage population^43^ and implicated ETA in the regulation of tumour-derived extracellular vesicle PD-L1^46^. Collectively, dual pharmacological inhibition of ETA and ETB has been shown to enhance responses to immune checkpoint inhibition across these preclinical settings^42,43,46^.

To test this hypothesis, we evaluated endothelin receptor blockade in combination with tumour-draining lymph node-derived adoptive T cell therapy (ACT) in the orthotopic 4T1 triple-negative breast cancer model and with anti-CD19 CAR T cell therapy in the A20 B-cell lymphoma model.

## RESULTS

### Endothelin receptor blockade enhances adoptive T cell therapy and, together with immune checkpoint inhibition, promotes durable tumour control

We first established an ACT model using the 4T1 triple-negative breast cancer orthotopic model. Isolated TDLN-derived T cells were activated and expanded *in vitro* using stimulatory antibody coated-beads and interleukin-2 (IL-2) (Fig. S1a-d). The seven-day expanded T cell end-product exhibited cytotoxic activity against 4T1 cells *in vitro* (Fig. S1d-f) and produced measurable but limited antitumour activity when administered as monotherapy *in vivo* (Fig. S2). These findings established a functional ACT product for subsequent combination therapy studies *in vivo*.

To determine whether inhibition of the endothelin receptor pathway could enhance the efficacy of ACT, mice bearing established 4T1 tumours were treated with TDLN-derived T cells either alone or in combination with bosentan-mediated endothelin receptor blockade (ERB), using the dual ETA/ETB antagonist bosentan and/or ICI (anti-PD-1 and anti-CTLA-4 blocking antibodies) (Fig. 1a). ACT monotherapy exhibited limited therapeutic activity, producing only a modest delay in tumour progression compared with untreated controls. Addition of ICI to ACT produced a moderate improvement in tumour control, whereas addition of ERB to ACT resulted in a more pronounced reduction in tumour growth. The greatest suppression of tumour progression was observed in mice receiving ACT, ERB and ICI, which maintained the lowest tumour volumes among the ACT-containing treatment groups throughout the observation period (Fig. 1a, Fig S3b-f). Consistent with previous reports^42,43,46^, ERB combined with ICI produced greater tumour control than either monotherapy alone (Fig. S3g-n). Importantly, the addition of ACT to ERB-ICI further enhanced tumour control, with ACT-ERB-ICI resulting in significantly lower tumour volumes at Day 28 compared with ERB-ICI (*P* = 0.0085). Body weight was largely maintained across treatment groups following completion of the treatment regimens, with no animals with paired measurements reaching the predefined 20% body-weight-loss humane endpoint (Fig. S3j).

**Fig. 1:**
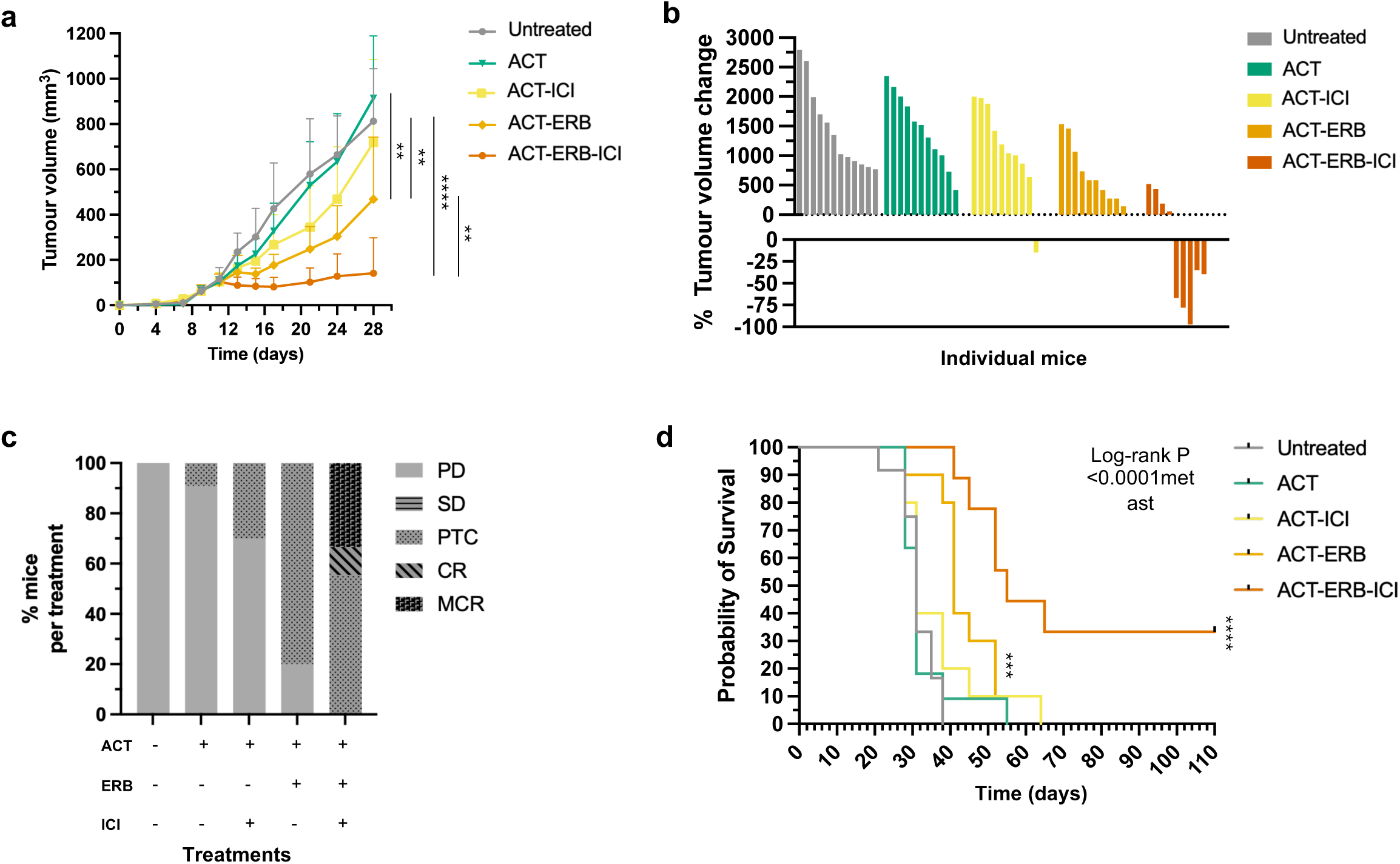
Endothelin receptor blockade enhances adoptive T-cell therapy and, together with immune checkpoint inhibition, promotes durable tumour control in the 4T1 cancer model. 4T1 tumour-bearing mice were treated with ACT alone (n = 11), ACT plus bosentan-mediated endothelin receptor blockade (ACT-ERB; n = 10), ACT plus immune checkpoint inhibition (ACT-ICI; n = 10), or the triple combination of ACT, endothelin receptor blockade and immune checkpoint inhibition (ACT-ERB-ICI; n = 9), and compared with untreated controls (n = 12). Treatment groups are colour-coded throughout the figure as follows: Control (grey), ACT (green), ACT-ICI (yellow), ACT-ERB (orange), and ACT-ERB-ICI (dark orange). **(a)** Tumour growth kinetics following treatment. Tumour volumes were measured longitudinally and are presented as mean ± SD. Bosentan-mediated endothelin receptor blockade (ERB) was initiated once tumours reached an average diameter of approximately 5 mm, which typically occurred around Day 9 after tumour inoculation. ACT was administered one day after initiation of ERB. ERB was administered daily thereafter, except on the day of ACT administration and ICI was initiated one day after ACT and followed by additional doses at three-day intervals, as shown in the treatment schema (Fig. S3a). Statistical comparisons were performed using two-tailed unpaired t-tests with Welch’s correction at Day 28. \*\**P* < 0.0064, \*\*\*\**P* < 0.0001 **(b)** Waterfall plot showing the percentage change in tumour volume for individual mice between ERB treatment initiation (Day 10) and completion of the treatment regimen (Day 28). Each bar represents an individual mouse. **(c)** Tumour response classification. Individual responses were classified using study-defined criteria based on tumour growth kinetics during treatment and longitudinal follow-up. Responses were classified as progressive disease (PD), stable disease (SD), partial tumour control (PTC), complete response (CR) or maintained complete response (MCR). Data are presented as the percentage of mice in each response category. Tumour control responses comprise PTC, CR and MCR. See Methods for the definition and evaluation of each response category. **(d)** Kaplan-Meier survival analysis of 4T1 tumour-bearing mice treated with ACT alone or in combination with bosentan-mediated endothelin receptor blockade (ERB) and/or immune checkpoint inhibition. Survival differed significantly among treatment groups (log-rank Mantel-Cox test, χ² = 29.74, df = 4, *P* < 0.0001). Pairwise comparisons demonstrated significantly improved survival for ACT-ERB versus ACT alone (*P* = 0.0283), ACT-ERB-ICI versus ACT alone (*P* = 0.0003), and ACT-ERB-ICI versus ACT-ERB (*P* = 0.0063).

Waterfall plot analysis of the percentage change in tumour volume between treatment initiation and treatment completion further illustrated the distribution of responses across treatment groups (Fig. 1B, S3l). ACT monotherapy resulted predominantly in continued tumour growth, while addition of ICI to ACT produced only a modest shift in tumour response. In contrast, addition of ERB to ACT shifted the response distribution towards substantially reduced tumour expansion. The greatest reductions in tumour volume were observed following triple-combination therapy with ACT, ERB and ICI, with several mice exhibiting tumour regression relative to pretreatment tumour size.

These differences were reflected in the overall tumour response classifications (Fig. 1c, S3m). Tumour responses were classified as progressive disease (PD), stable disease (SD), partial tumour control (PTC), complete response (CR) or maintained complete response (MCR) based on tumour growth kinetics during treatment and longitudinal follow-up, according to the study-defined criteria described in Methods. PTC, CR and MCR were collectively considered tumour control responses, reflecting treatment-associated tumour control ranging from reduced tumour progression to complete and maintained tumour regression. ACT monotherapy resulted predominantly in progressive disease, with only a single PTC observed (1/11 mice) (Fig. 1c). Addition of ICI to ACT modestly increased the frequency of tumour control responses, from 1 of 11 mice receiving ACT alone to 3 of 10 mice receiving ACT-ICI, with all tumour control responses in both groups classified as PTC. Addition of ERB to ACT resulted in tumour control responses in the majority of treated animals (8/10 mice, all PTC). In contrast, mice receiving ACT-ERB-ICI exhibited no progressive disease and included both complete and maintained complete responses. MCRs were observed exclusively in this triple-combination therapy group, with 3 of 9 mice achieving complete tumour regression without recurrence throughout the 110-day observation period.

Overall survival analysis mirrored the response classification data (Fig. 1d). ACT monotherapy did not significantly improve survival compared with untreated controls, and the addition of ICI to ACT did not significantly extend survival relative to ACT alone. In contrast, the addition of ERB to ACT significantly prolonged survival compared with ACT monotherapy (log-rank *P* = 0.0283). Consistent with previous reports^42,43^, ERB combined with ICI also improved survival relative to either monotherapy alone (Fig. S3n). However, the greatest survival benefit was observed in mice receiving ACT-ERB-ICI, which significantly prolonged survival compared with ACT alone (*P* = 0.0003), ACT-ERB (*P* = 0.0063) and ERB-ICI (*P* = 0.0165). Consistent with the response classification analysis, durable long-term survival was observed only in the triple-combination group, with 33.3% of animals remaining alive and tumour-free at study completion.

Together, these findings demonstrate that ERB markedly enhances the antitumour activity of ACT by increasing the frequency of tumour control responses. The addition of ICI to ACT-ERB promotes the emergence of maintained complete responses and provides the greatest durability of tumour control. Importantly, ACT also provides additional therapeutic benefit beyond combined ERB-ICI, with the triple combination producing significantly greater tumour control and prolonged survival. Notably, maintained complete responses and long-term tumour-free survival were observed exclusively following triple-combination therapy.

### Maintained complete responses following combined ACT therapy, endothelin receptor blockade and immune checkpoint inhibition are associated with resistance to tumour rechallenge

To determine whether durable antitumour protection had been established following successful therapy, mice achieving maintained complete responses (MCRs) after treatment with ACT-ERB-ICI were rechallenged with 4T1 tumour cells at Day 110 following primary tumour inoculation. Tumour cells were implanted into the contralateral mammary fat pad, with treatment-naïve mice of the same age challenged in parallel as controls (Fig. 2a-b).

**Fig. 2:**
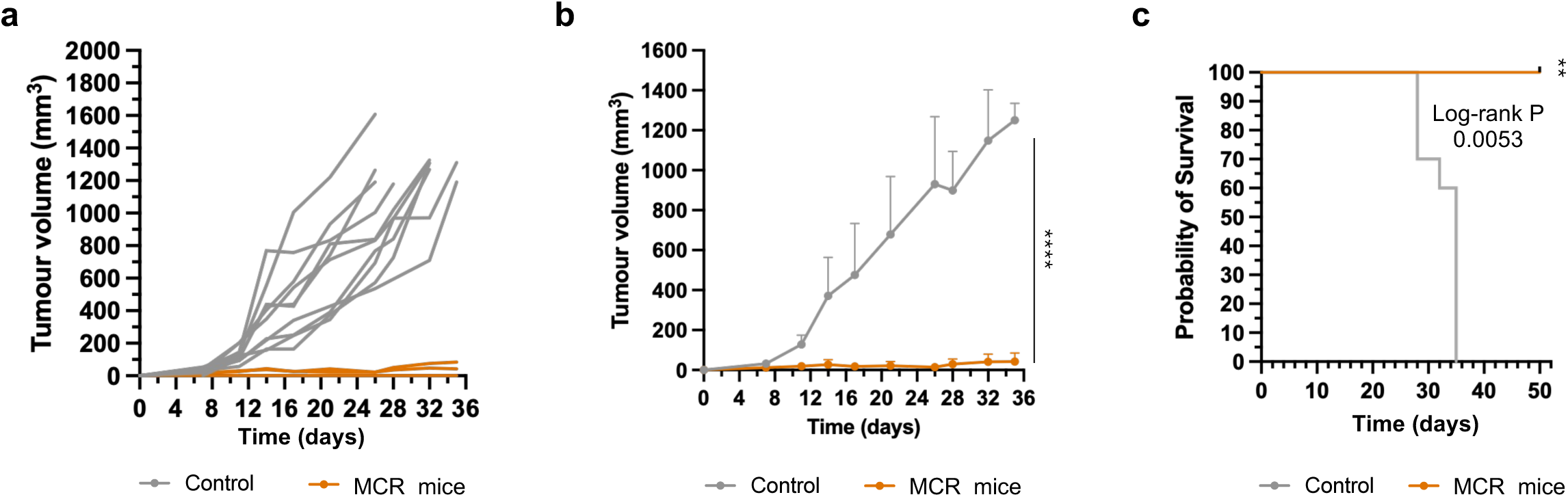
Maintained complete responses following combined ACT, endothelin receptor blockade and immune checkpoint inhibition are associated with resistance to tumour rechallenge. Mice achieving maintained complete responses (MCRs) following ACT-ERB-ICI treatment in the primary 4T1 tumour experiment were rechallenged with the same number of 4T1 tumour cells at Day 110 following primary tumour inoculation. Tumour cells were implanted into the contralateral mammary fat pad and compared with treatment-naïve control mice challenged in parallel. Control mice are shown in grey and rechallenged MCR mice in dark orange. **(a)** Individual tumour growth curves following tumour rechallenge. Each line represents an individual mouse. Naïve control mice (n = 10) developed progressively growing tumours, whereas MCR mice (n = 3) either completely rejected the rechallenge or exhibited only limited tumour growth that remained controlled throughout the observation period. **(b)** Mean tumour growth kinetics following tumour inoculation in the two groups described above. Tumour volumes were measured longitudinally and are presented as mean ± SD. Statistical comparisons were performed using two-tailed unpaired t-tests with Welch’s correction at Day 32, when six of ten control mice and all three rechallenged MCR mice remained evaluable. \*\*\*\**P* < 0.0001 **(c)** Kaplan-Meier survival analysis following tumour inoculation in the two groups described above. Survival differed significantly between treatment-naïve control mice and rechallenged MCR mice (log-rank Mantel-Cox test, χ² = 7.760, df = 1, *P* = 0.0053). All rechallenged MCR mice survived until study completion, whereas no control animals remained alive beyond Day 35.

All naïve control mice developed progressively growing tumours following challenge and reached endpoint within 35 days. In contrast, the aforementioned MCR mice exhibited marked resistance to tumour re-establishment. One of three rechallenged MCR mice completely rejected tumour growth, while the remaining two mice displayed only minimal tumour expansion that remained controlled throughout the study period. Consistent with this protection, all rechallenged MCR mice survived until study completion, whereas no control animals remained alive beyond Day 35 (log-rank Mantel-Cox test, χ² = 7.760, *P* = 0.0053).

Taken together, these findings demonstrate substantial protection against tumour development following 4T1 rechallenge in mice that achieved maintained complete responses to ACT-ERB-ICI therapy. Notably, all rechallenged MCR mice either completely rejected tumour growth or maintained minimal tumour expansion throughout the study period, whereas all treatment-naïve controls developed progressively growing tumours and reached endpoint, consistent with durable antitumour immune protection.

### Treatment-associated changes in intratumoural T cell abundance

Given that the primary objective of this study was to identify strategies for enhancing ACT efficacy, we next examined treatment-associated changes within the intratumoural T cell compartment. Tumours harvested following completion of the treatment regimens were therefore analysed by immunofluorescence staining for CD8⁺ T cells, granzyme B-expressing CD8⁺ T cells (GzmB⁺CD8⁺), and CD4⁺ T cells (Fig. 3a-b).

**Fig. 3:**
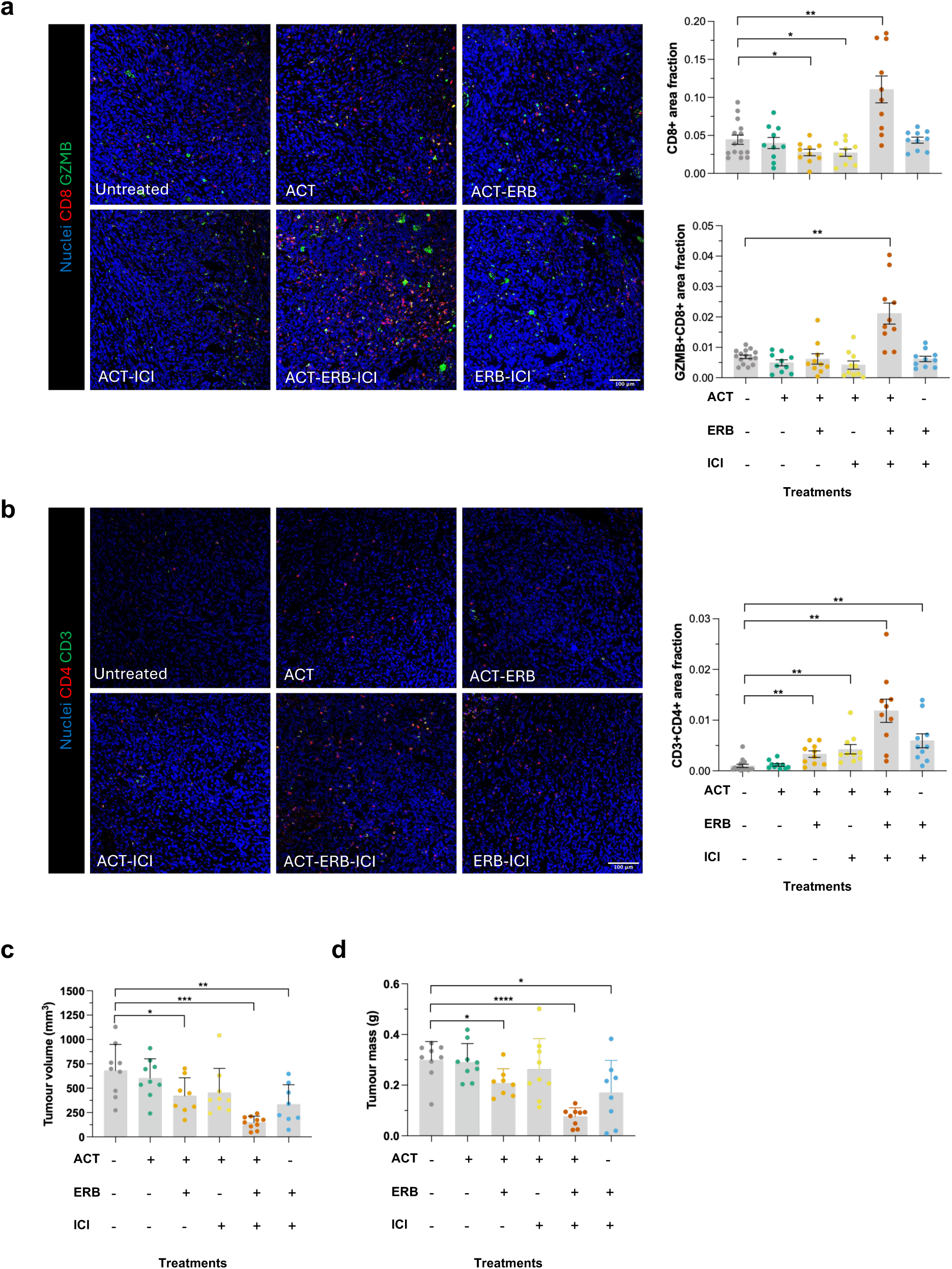
Treatment-associated changes in intratumoral T-cell abundance. Tumours were harvested following completion of the treatment regimens and analysed by immunofluorescence staining to quantify intratumoral CD8⁺ T cells, granzyme B-expressing CD8⁺ T cells and CD4⁺ T cells. Treatment groups are colour-coded throughout the figure as follows: Untreated (grey), ACT (green), bosentan-mediated endothelin receptor blockade plus immune checkpoint inhibition (ERB-ICI; light blue), ACT plus ERB (ACT-ERB; yellow-orange), ACT plus ICI (ACT-ICI; yellow) and ACT plus ERB and ICI (ACT-ERB-ICI; dark orange). **(a)** Representative immunofluorescence images and quantification of total CD8⁺ T cells and granzyme B-expressing CD8⁺ T cells (GzmB⁺CD8⁺) within tumour sections. CD8 is shown in red, granzyme B in green, and nuclei (DAPI) in blue. **(b)** Representative immunofluorescence images and quantification of intratumoral CD4⁺ T cells (CD3⁺CD4⁺). CD4 is shown in red, CD3 in green and nuclei (DAPI) in blue. For immunofluorescence analyses, five fields of view were analysed per tumour from 2-3 tumours per treatment group; individual symbols represent image fields. Bar graphs represent mean ± SD. Statistical comparisons shown on the graphs were performed using two-tailed unpaired t-tests with Welch’s correction. \**P* ≤ 0.05, \*\**P* ≤ 0.01, \*\*\**P* ≤ 0.001, \*\*\*\**P* ≤ 0.0001. Scale bars, 100 µm. **(c)** Tumour volume measured on the day of tumour harvest. **(d)** Tumour mass measured on the day of tumour harvest.

Total CD8⁺ T cell abundance was significantly increased only in the ACT-ERB-ICI treatment group relative to control, whereas significant reductions were observed in the ACT-ERB and ACT-ICI groups, with no significant differences detected in ACT or ERB-ICI groups. Consistent with this, GzmB⁺CD8⁺ T cell abundance was also significantly increased only in the ACT-ERB-ICI group, with no significant changes observed in the remaining treatment groups.

In contrast, CD4⁺ T cell abundance was significantly increased in all treatment groups except ACT, with increases of approximately 3.3-fold in ACT-ERB, 4.3-fold in ACT-ICI, 6.0-fold in ERB-ICI, and 12.0-fold in ACT-ERB-ICI relative to control.

Tumour burden was assessed at the same endpoint to examine whether changes in intratumoural T cell populations were associated with therapeutic efficacy (Fig. 3c-d). Tumour volume was significantly reduced relative to control in the ACT-ERB (*P ≤ 0.05), ERB-ICI (**P ≤ 0.01), and ACT-ERB-ICI (***P ≤ 0.001) groups. Similarly, tumour mass was significantly reduced in the ACT-ERB (*P ≤ 0.05), ERB-ICI (*P ≤ 0.05), and ACT-ERB-ICI (****P ≤ 0.0001) groups. The greatest reductions in both tumour volume and tumour mass were observed in the ACT-ERB-ICI group.

Taken together, these findings show that while increased CD4⁺ T cell abundance occurred across multiple treatment groups, ACT-ERB-ICI was uniquely associated with concurrent increases in CD4⁺ T cells, total CD8⁺ T cells and GzmB⁺CD8⁺ T cells, alongside the greatest reduction in tumour burden.

### High-dimensional phenotypic profiling identifies multiple intratumoral T cell states associated with tumour control

Tumour-infiltrating T cell populations are highly heterogeneous and can comprise functionally distinct subsets with differing capacities for tumour control. We therefore performed high-dimensional phenotypic profiling to investigate intratumoral T cell heterogeneity and determine whether specific T cell states were associated with tumour volume.

Spectral flow cytometric data were analysed using t-distributed stochastic neighbour embedding (tSNE) dimensionality reduction followed by Phenograph clustering based on expression of 12 surface markers. This approach identified 18 phenotypically distinct T cell populations within the tumour microenvironment (Fig. 4a-b, Tables S1-2). Correlation analysis between cluster frequencies and tumour volume at harvest identified distinct CD4⁺, CD8⁺ and double-negative T cell populations associated either positively or negatively with tumour volume.

**Fig. 4:**
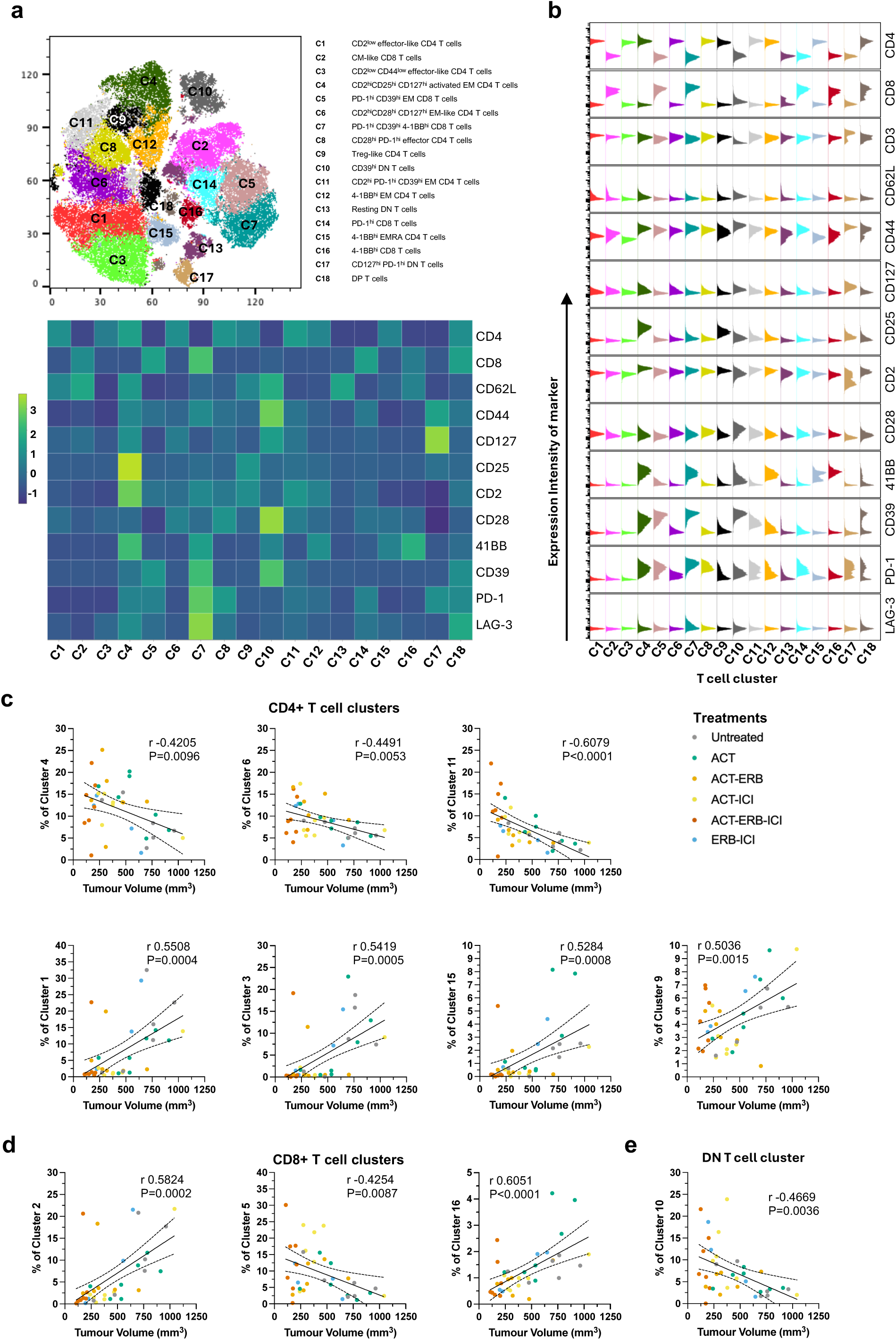
High-dimensional phenotypic profiling identifies multiple intratumoral T-cell states associated with tumour control. Tumours harvested at the same endpoint as the histology cohort were analysed by spectral flow cytometry following gating on live CD45⁺CD49b^-^B220^-^CD3⁺ tumour infiltrating T cells. High-dimensional phenotypic profiling was performed using t-SNE dimensionality reduction followed by Phenograph clustering as described in Methods. **(a)** t-SNE projection of intratumoral T cells coloured according to Phenograph cluster assignment together with a Cluster Explorer heatmap showing normalised mean marker expression across the 18 identified T cell clusters. **(b)** Histograms showing marker expression profiles for each Phenograph-defined cluster used for phenotypic annotation of intratumoral T cell populations. Correlation analyses between cluster frequency and tumour volume were performed using Pearson correlation across pooled tumours from all treatment groups. Each data point represents an individual mouse and is colour-coded according to treatment group assignment using the same colour scheme as in previous figures. **(c)** Correlation analyses between the frequencies of CD4⁺ T cell clusters and tumour volume at harvest. **(d)** Correlation analyses between the frequencies of CD8⁺ T cell clusters and tumour volume at harvest. **(e)** Correlation analyses between the frequencies of double-negative (CD4⁻CD8⁻) T cell clusters and tumour volume at harvest. Regression lines are shown with 95% confidence intervals. Pearson correlation coefficients (r) and corresponding *P* values are displayed within each panel. Only statistically significant correlations are shown.

Among CD4⁺ T cells, three phenotypically distinct clusters (C4, C6 and C11) were significantly associated with lower tumour volume (Fig. 4c) (C4: *r* = −0.42, R² = 0.18, *P* = 0.0096; C6: *r* = −0.45, R² = 0.20, *P* = 0.0053; C11: *r* = −0.61, R² = 0.37, *P* < 0.0001). Although these populations differed in their activation-marker profiles, all displayed features consistent with antigen-experienced and/or effector-memory T cell states. Cluster 4 displayed the highest CD2 expression of all identified clusters and was characterised by concurrent expression of CD25, CD127, 4-1BB, PD-1 and CD39. Notably, despite expressing PD-1 and CD39, cluster 4 also ranked among the five highest CD127-expressing populations. Together with its high CD25 and 4-1BB expression, this phenotype is consistent with a highly activated, antigen-experienced effector-memory CD4⁺ T cell population despite expression of markers commonly associated with chronic antigen stimulation. Cluster 6 was characterised by high CD2, CD28 and CD127 expression together with relatively low PD-1, CD39, CD25 and absent 4-1BB expression, consistent with a CD28^hi^CD127^hi^ effector-memory-like CD4⁺ T cell phenotype. Cluster 11 combined high CD2, PD-1 and CD39 expression with relatively low CD25 and minimal 4-1BB expression, defining a phenotypically distinct antigen-experienced effector-memory CD4⁺ population. Notably, cluster 11 exhibited the strongest inverse association with tumour volume among all identified T cell populations.

In contrast, four phenotypically distinct CD4⁺ clusters (C1, C3, C9 and C15) were positively associated with tumour volume (Fig. 4d) (C1: *r* = 0.55, R² = 0.30, *P* = 0.0004; C3: *r* = 0.54, R² = 0.29, *P* = 0.0005; C9: *r* = 0.50, R² = 0.25, *P* = 0.0015; C15: *r* = 0.53, R² = 0.28, *P* = 0.0008). Clusters 1 and 3 expressed low CD62L and CD44, while cluster 15 combined low CD62L with higher CD44 and moderate-to-high 4-1BB. Cluster 9, annotated as a regulatory T cell-like population (CD25^hi/int^CD127^low^), showed a positive association with tumour volume. With the exception of C9, these populations exhibited lower CD2 expression than the tumour-control-associated CD4⁺ clusters.

Within the CD8⁺ compartment, one cluster (C5) was significantly associated with lower tumour volume, whereas two clusters (C2 and C16) were positively associated with tumour volume (Fig. 4d). Cluster 5 (*r* = −0.43, R² = 0.18, *P* = 0.0087) exhibited a PD-1^hi^CD39^hi^CD44^hi^CD62L^low^CD2i^nt/hi^4-1BB^low^ phenotype consistent with an activated antigen-experienced effector-memory CD8⁺ T cell population. In contrast, C2 and C16 (C2: *r* = 0.58, R² = 0.34, *P* = 0.0002; C16: *r* = 0.61, R² = 0.37, *P* < 0.0001) expressed low PD-1, CD39 and CD2; C16, however, expressed high 4-1BB.

Among double-negative (CD4⁻CD8⁻) T cells, cluster 10 was significantly associated with lower tumour volume (*r* = −0.47, R² = 0.22, *P* = 0.0036) (Fig. 4e) and exhibited a highly activated phenotype characterised by very high CD39, CD44, CD28 and 4-1BB expression.

Together, these findings demonstrate that reduced tumour volume was associated with several phenotypically distinct intratumoral T cell populations spanning the CD4⁺, CD8⁺ and double-negative compartments (C4, C5, C6 and C10-11). Despite their diverse activation and differentiation phenotypes, these tumour-control-associated populations consistently exhibited higher CD2 expression than populations positively associated with tumour volume, identifying elevated CD2 expression as a common feature shared across multiple phenotypically distinct T cell states associated with effective tumour control. High-dimensional phenotypic profiling therefore complemented conventional measures of intratumoral T cell abundance by resolving biologically meaningful heterogeneity within the tumour T cell compartment and identifying multiple phenotypically distinct T cell states associated with reduced tumour volume.

### Endothelin receptor blockade enhances CAR T cell therapy and, together with immune checkpoint inhibition, promotes durable tumour control

Having established that endothelin receptor blockade enhances the efficacy of adoptively transferred non-engineered T cells, we next investigated whether this approach could similarly improve engineered CAR T cell therapy. To address this, we evaluated anti-CD19 CAR T cell therapy (CAR19) in the A20 B cell lymphoma model, either alone or in combination with ERB with or without ICI (Fig. 5 and Fig. S6). Prior to in vivo evaluation, transduction efficiency was assessed by flow cytometric detection of the Thy1.1 marker encoded together with the CAR construct (Fig. S6b), and the functional activity of CAR19 T cells against A20 cells was assessed in four independent in vitro cytotoxicity assays. CAR19 T cells exhibited significantly greater cytotoxicity against A20 cells than against antigen-mismatched Raji cells at both 1:1 (*P* = 0.0013) and 5:1 (*P* = 0.0003) effector-to-target ratios, with A20 cytotoxicity increasing further at the higher effector-to-target ratio (*P* = 0.0036) (Fig. S6c).

**Fig. 5.**
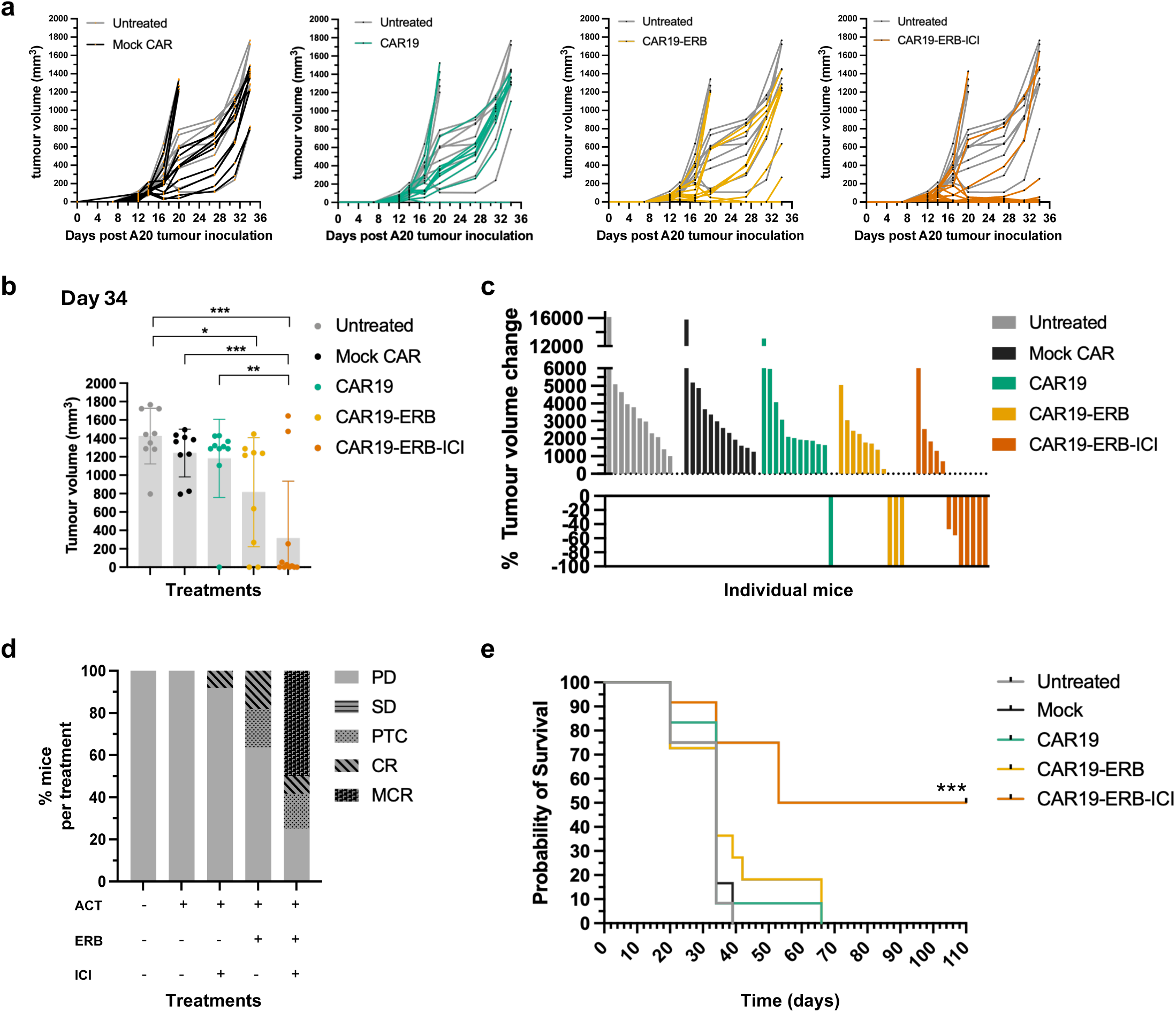
Endothelin receptor blockade enhances CAR T-cell therapy and, together with immune checkpoint inhibition, promotes durable tumour control. **(a)** Individual tumour growth trajectories of A20 tumour-bearing mice treated with control, mock CAR T cells, anti-CD19 CAR T cells alone, anti-CD19 CAR T cells combined with bosentan-mediated endothelin receptor blockade (CAR19-ERB), or anti-CD19 CAR T cells combined with endothelin receptor blockade and immune checkpoint inhibition (CAR19-ERB-ICI; anti-PD-1 and anti-CTLA-4). Each line represents an individual mouse. **(b)** Tumour volumes measured at Day 34 following tumour inoculation. Each symbol represents an individual mouse. Statistical comparisons between groups were performed using two-tailed unpaired t-tests with Welch’s correction. Significant differences were observed for CAR19-ERB versus control (\**P* = 0.0177), CAR19-ERB-ICI versus control (\*\*\**P* = 0.0001), CAR19-ERB-ICI versus mock CAR (\*\*\**P* = 0.0005), and CAR19-ERB-ICI versus CAR19 (\*\**P* = 0.0015). **(c)** Waterfall plot showing the percentage change in tumour volume between the pretreatment timepoint and completion of the treatment regimen for individual mice. Negative values indicate tumour regression, with −100% representing complete tumour regression. **(d)** Tumour response classification. Individual responses were classified using study-defined criteria based on tumour growth kinetics during treatment and longitudinal follow-up. Responses were classified as progressive disease (PD), stable disease (SD), partial tumour control (PTC), complete response (CR) or maintained complete response (MCR). Data are presented as the percentage of mice in each response category. Tumour control responses comprise PTC, CR and MCR. See Methods for the definition and evaluation of each response category. **(e)** Kaplan-Meier survival analysis of A20 tumour-bearing mice treated with mock CAR T cells, anti-CD19 CAR T cells alone, CAR19-ERB, or CAR19-ERB-ICI. Statistical significance was assessed using the log-rank (Mantel-Cox) test.

Individual tumour growth trajectories showed that CAR19 monotherapy did not markedly improve tumour control compared with mock CAR T cell treatment. In contrast, combining CAR19 with ERB delayed tumour progression and induced tumour regression in a subset of mice, while the addition of ICI to CAR19-ERB produced the most pronounced tumour control (Fig. 5a-b, Fig S6d). Consistent with the individual tumour growth trajectories, analysis at the common Day 34 endpoint, before substantial attrition resulting from humane endpoints, revealed a broader distribution of tumour responses in mice receiving CAR19 therapy combined with ERB, with the lowest tumour volumes observed in the triple-combination group, CAR19-ERB-ICI (Fig.5b, Fig. S6d).

Waterfall plot analysis comparing the percentage change in tumour volume between the pretreatment timepoint and completion of the treatment regimen further illustrated the enhanced antitumour activity achieved by combining CAR19 with ERB, with the greatest percentage reductions in tumour volume observed in mice receiving CAR19-ERB-ICI (Fig. 5c).

The overall tumour response classifications further demonstrated that addition of ERB substantially increased the frequency of tumour control responses, including complete tumour regressions, compared with CAR19 alone (Fig. 5d). Although CAR19-ERB induced complete responses, these were not consistently maintained, whereas the further addition of ICI increased the frequency of tumour control responses and promoted the emergence of maintained complete responses, indicating enhanced durability of tumour control. MCRs were observed exclusively in the CAR19-ERB-ICI group, with 6 of 12 mice achieving complete tumour regression without recurrence throughout the 110-day observation period. Consistent with these response profiles, CAR19-ERB-ICI produced the greatest survival benefit and was the only regimen associated with long-term survivors at the end of the study period (Fig. 5e).

Together, these data demonstrate that ERB enhances otherwise limited CAR19 activity by increasing the frequency of tumour control responses, including tumour regressions, while the addition of ICI to CAR19-ERB further increases the frequency of tumour control responses and promotes the emergence of maintained complete responses.

### Maintained complete responses following combined CAR T cell therapy, endothelin receptor blockade and immune checkpoint inhibition are associated with resistance to tumour rechallenge

To determine whether maintained complete responses that were generated only following CAR19-ERB-ICI therapy were associated with long-term protection against tumour rechallenge, maintained complete responder mice were rechallenged with A20 tumour cells. Treatment-naïve mice challenged in parallel served as controls.

Following rechallenge, all treatment-naïve control mice developed progressively growing tumours. In contrast, four of six maintained complete responder mice remained tumour-free throughout the observation period, while tumour outgrowth was observed in only two animals (Fig. 6a). Consistent with these tumour growth patterns, rechallenged maintained complete responder mice exhibited significantly prolonged tumour-free survival compared with treatment-naïve controls (Fig. 6b).

**Fig. 6:**
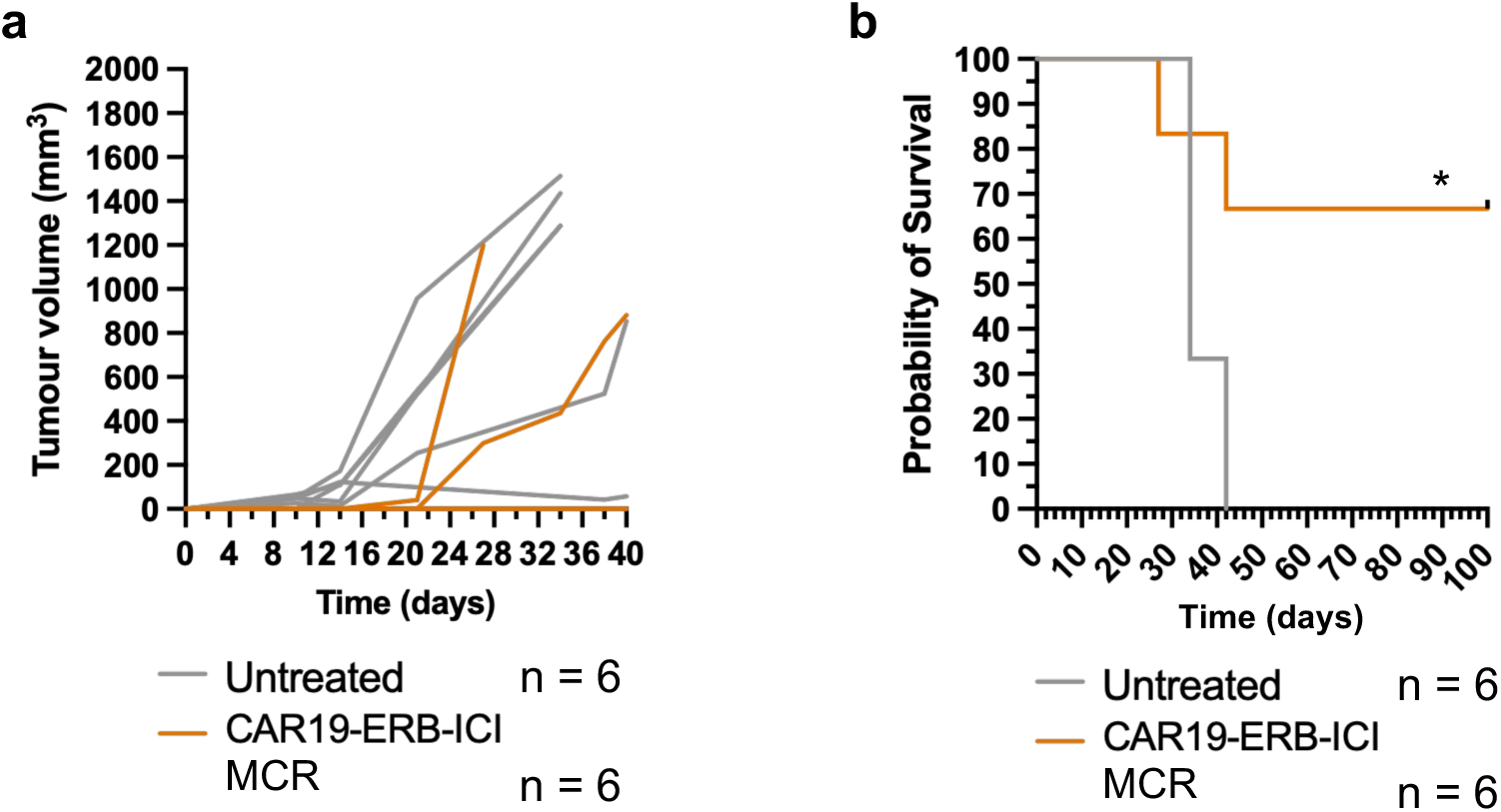
Maintained complete responses following combined CAR T-cell therapy, endothelin receptor blockade and immune checkpoint inhibition are associated with resistance to tumour rechallenge. **(a)** Tumour growth following rechallenge of maintained complete responder (MCR) mice previously treated with anti-CD19 CAR T cells, bosentan-mediated endothelin receptor blockade and immune checkpoint inhibition (CAR19-ERB-ICI), and treatment-naïve control mice with A20 lymphoma cells injected subcutaneously into the contralateral mammary fat pad. Rechallenge was performed 110 days after primary tumour inoculation. Each line represents an individual mouse. **(b)** Kaplan-Meier analysis of tumour-free survival following A20 tumour rechallenge. Statistical significance was assessed using the log-rank (Mantel-Cox) test (χ² = 4.479, *P* = 0.0343). Four of six maintained complete responder mice remained tumour-free following rechallenge, whereas all treatment-naïve control mice developed progressive tumours.

Taken together, these findings demonstrate substantial protection against tumour development following A20 rechallenge in mice that achieved durable responses to combined CAR T cell therapy, endothelin receptor blockade and immune checkpoint inhibition. Notably, most MCR mice remained tumour-free following rechallenge 110 days after primary tumour inoculation, whereas all treatment-naïve controls developed progressively growing tumours, consistent with durable antitumour immune protection.

## DISCUSSION

Adoptive T cell therapies have transformed the treatment of selected haematological malignancies, yet their efficacy remains limited in many tumour settings by barriers that restrict the generation, maintenance and durability of effective antitumour responses by adoptively transferred T cells. In this study, we investigated whether pharmacological inhibition of the endothelin receptor pathway could enhance the efficacy of T cell-based immunotherapy across two distinct platforms: non-engineered ACT in the orthotopic 4T1 triple-negative breast cancer model and anti-CD19 CAR T cell therapy in the A20 B-cell lymphoma model. In both settings, ERB significantly increased the frequency of tumour control responses achieved with T cell therapy, while the addition of ICI to the combination of ERB and T cell therapy increased the frequency of tumour control responses and produced the most durable tumour control. Therapeutic responses were associated with multiple intratumoral T cell populations identified by high-dimensional immune profiling, providing new insights into the phenotypic T cell states associated with effective tumour control. Collectively, these findings suggest that pharmacological inhibition of the endothelin receptor pathway may represent a promising rational strategy to address two of the major challenges facing T cell-based cellular immunotherapy: increasing the frequency of tumour control responses and enhancing the durability of tumour control.

An important observation from this study is that ERB enhanced the efficacy of both non-engineered ACT and engineered CAR T cell therapy by increasing the frequency of tumour control responses. Although the underlying mechanisms remain to be defined, these findings are consistent with the hypothesis that ERB alleviates biological barriers that limit effective antitumour responses mediated by adoptively transferred T cells. The reproducibility of this therapeutic benefit across two distinct tumour models and T cell therapy platforms suggests that this strategy may have broader applicability, although further validation across additional tumour types will be important.

One of the most notable findings of this study was the complementary contribution of ERB and ICI to therapeutic efficacy. Although both interventions enhanced the therapeutic activity of adoptively transferred T cells, their relative effects differed. ERB was associated with a marked increase in the frequency of tumour control responses, whereas the addition of ICI to the combination of ERB and T-cell therapy further increased the frequency of tumour control responses and exerted a particularly pronounced effect on the durability of tumour control, promoting the emergence of maintained complete responses. These observations are consistent with the possibility that ERB and ICI act through complementary biological processes that together improve both the frequency and durability of responses to adoptively transferred T-cell therapies. The observation of this pattern independently in both ACT and CAR T cell settings further supports investigation of how these interventions may influence distinct but complementary biological barriers limiting transferred T-cell efficacy.

Although generation of initial tumour regression remains an important objective of cellular immunotherapy, the maintenance of long-term tumour control represents one of the major challenges limiting the efficacy of both ACT and CAR T cell therapies. In both models examined here, ERB increased the frequency of tumour control responses, whereas the addition of ICI to T cell therapy combined with ERB was most clearly associated with the emergence of maintained complete responses and durable tumour control. Particularly notable was the observation that the majority of maintained complete responder mice from the 4T1 ACT and A20 CAR T cell studies resisted 4T1 and A20 tumour rechallenge, respectively, when rechallenged 110 days after primary tumour inoculation, whereas all treatment-naïve controls developed progressive tumours. Although the protection mechanisms were not investigated directly, rejection of a secondary tumour challenge is consistent with the development of durable antitumour immune memory^30^. These findings further argue that the triple-combination strategy not only dramatically increases the frequency of tumour control responses but also promotes the establishment of long-term immune control, an important objective for preventing disease recurrence following cellular immunotherapy.

A further noteworthy aspect of the study is that therapeutic responses were generated without host lymphodepleting conditioning in either experimental system. Lymphodepletion preconditioning of the host is commonly incorporated into adoptive cellular therapy protocols^47^ to enhance transferred T cell activity, in part by increasing the availability of homeostatic cytokines through the removal of endogenous cellular cytokine sinks^48^. Here, both ACT and CAR T cell therapies exhibited limited efficacy when administered alone in non-conditioned hosts, yet tumour control responses emerged following the addition of ERB. Although our study was not designed to determine whether ERB can substitute for lymphodepleting conditioning, these findings raise the possibility that ERB may partially overcome barriers restricting effective T cell-mediated tumour control in non-conditioned settings. This possibility warrants further investigation, including studies directly comparing endothelin receptor blockade with conventional lymphodepleting conditioning strategies.

The therapeutic benefit observed in the 4T1 ACT model was accompanied by marked alterations in the intratumoral T cell compartment. Immunofluorescence analyses demonstrated increased intratumoral CD4⁺ T cell abundance together with increased numbers of CD8⁺ and granzyme B-expressing CD8⁺ T cells, particularly following triple-combination therapy. These findings established a clear association between enhanced intratumoral representation of both helper and cytotoxic T cell populations and tumour control. However, while these analyses provided important information regarding T cell abundance, tumour-infiltrating T cells are phenotypically heterogeneous and may differ substantially in their functional capacity. We therefore performed high-dimensional phenotypic profiling to determine whether specific intratumoral T cell states, beyond bulk measures of T cell abundance, were associated with therapeutic outcome. Phenograph clustering revealed a considerably more complex intratumoral T cell landscape in which multiple CD4⁺, CD8⁺ and double-negative T cell populations were differentially associated with tumour volume. Therapeutic efficacy may therefore depend not only on T cell abundance but also on the balance between phenotypically distinct states associated with tumour control or tumour persistence.

The tumour-control-associated T cell populations were phenotypically diverse. Several expressed PD-1 and/or CD39, markers frequently associated with chronic antigen stimulation and T cell dysfunction^49,50^. However, these same populations also displayed features consistent with activation, memory formation or co-stimulatory competence, including high expression of CD2, CD127 and/or 4-1BB. These findings argue against interpreting PD-1 or CD39 expression in isolation as evidence of dysfunction and instead support the emerging view that these molecules can occur within tumour-reactive or antigen-experienced T-cell populations^51,52^, whose functional state should be considered within their broader phenotypic context. In our setting, the association of PD-1- and CD39-expressing populations with tumour control further raises the possibility that these molecules may mark tumour-engaged or antigen-experienced T cells with retained functional potential. Consistent with this interpretation, the strongest inverse correlation with tumour burden was observed in a CD2-high population co-expressing PD-1 and CD39, showing that PD-1 and CD39 expression can coexist within a T-cell population associated with effective tumour control.

Perhaps the most striking observation from the high-dimensional analysis was that, despite their marked phenotypic diversity, tumour-control-associated T cell populations consistently exhibited high CD2 expression compared with populations positively associated with tumour volume. This recurring association emerged independently across multiple CD4⁺, CD8⁺ and double-negative T cell populations identified through an unbiased analysis, rather than being confined to a single phenotypic lineage or differentiation state. Although additional markers associated with activation or co-stimulatory fitness, including CD28, CD127 and 4-1BB, were enriched within individual tumour-control-associated T cell populations, none showed the consistency observed for high CD2 expression across the full spectrum of favourable T cell populations. High CD2 expression may therefore represent a broader feature of intratumoural T cell populations associated with effective tumour control.

Activated T cells can upregulate CD2 expression, raising the possibility that high CD2 expression may partly reflect prior antigenic stimulation. However, the association observed here is unlikely to be explained solely by activation status. While some tumour-control-associated T cell populations expressed high levels of conventional activation markers such as CD25 or 4-1BB, others did not, yet high CD2 remained a recurring feature across these phenotypically distinct populations. CD2 may therefore reflect broader aspects of T cell fitness, tumour engagement or functional competence that extend beyond simple activation status.

The recurring emergence of high CD2 across phenotypically diverse tumour-control-associated T cell populations was particularly intriguing because it arose from an unbiased high-dimensional analysis rather than from a predefined hypothesis. Its potential biological significance is supported by growing evidence implicating the CD2 axis in productive antitumour T cell immunity beyond its classical role in T cell activation^53–56^. In our previous work, we demonstrated that the strength of CD2 co-stimulation depends on the extent of CD2 engagement by its ligand CD58, which is determined by both CD2 expression levels and CD58 availability^57^. We further showed that the levels of CD2 expression regulate formation of the novel CD2 corolla at the immunological synapse, a specialised signalling domain that amplifies proximal T cell receptor signalling and promotes downstream T cell activation^57^. We subsequently showed that variation in CD2 co-stimulation strength influences T cell proliferation and effector cytokine production^58^.

In our previous studies, CD8⁺CD3⁺ tumour-infiltrating lymphocytes (TILs) from untreated patients with colorectal, ovarian and endometrial cancers exhibited lower CD2 expression than matched peripheral blood CD8⁺ T-cell subsets^57^. Similarly, CD8⁺CD3⁺ TILs from patients with meningioma and glioblastoma exhibited reduced CD2 expression relative to their matched peripheral blood counterparts^58^. In these brain tumours, CD4⁺CD3⁺ TILs also exhibited reduced CD2 expression relative to matched peripheral blood CD4⁺ T-cell subsets^58^. These findings suggest that diminished CD2 expression may be a recurring feature of tumour-infiltrating T cells across multiple tumour types. Other studies have demonstrated that the CD2-CD58 axis directly influences T cell-mediated tumour control in xenograft and partially humanised mouse^55^ cancer models by regulating T cell activation and migration within the tumour microenvironment. More recently, engineered enhancement of CD2 expression has been shown to improve CAR T cell efficacy, strengthen immunological synapse formation, promote persistence and attenuate exhaustion-associated programmes in preclinical cancer models^59,60^. Together, these observations suggest that the levels of CD2 expression represent both a biomarker of favourable T cell states and a biologically relevant pathway with therapeutic potential. Although CD2 was not directly manipulated here, the independent emergence of high CD2 expression across multiple tumour-control-associated T cell populations, together with the growing body of evidence implicating the CD2 axis in productive antitumour T cell immunity, identifies it as an attractive candidate for future mechanistic investigation. Elucidating how ERB enhances the antitumour activity of adoptively transferred tumour-reactive T cells, with or without ICI, and how these interventions influence the generation and maintenance of favourable intratumoral T cell states therefore represent important priorities for future investigation.

Given the increasingly recognised pleiotropic effects of endothelin signalling on tumour biology and host immunity, ERB-mediated enhancement of adoptively transferred T cell therapy is likely to involve multiple, non-mutually exclusive mechanisms. These may include modulation of tumour vascular function^61,62^ and stromal remodelling within the tumour microenvironment^42,63^, altered immune-cell trafficking and/or function^23,42^, modulation of host immune-cell populations or their function^43^. Whether endothelin receptor blockade also acts directly on transferred T cells remains to be determined. Defining the relative contribution of these mechanisms, including their influence on intratumoral T cell states associated with tumour control, will be important for guiding the rational optimisation of future combination therapies.

The applicability of the triple-combination strategy may also extend to settings in which treatment is initiated following surgical resection of the primary tumour. In an exploratory post-surgical experiment, mice bearing established 4T1 tumours underwent surgical primary tumour resection before receiving the triple-combination therapy (ACT-ERB-ICI). Post-operative triple-combination therapy significantly prolonged survival compared with surgery alone (Fig. S7; log-rank *P* = 0.0398), with 4 of 10 treated mice remaining alive at Day 100 compared with 1 of 8 evaluable controls. Although local tumour recurrence contributed to endpoint determination in some animals, necropsy frequently revealed macroscopic metastatic disease, predominantly in the lungs, indicating that post-surgical disease progression often involved metastatic outgrowth, either alone or together with local tumour recurrence. As this exploratory study included only surgery-alone and triple-combination groups, it cannot define the contribution of individual components. Nevertheless, these findings provide preliminary evidence that initiation of the triple-combination strategy following surgical resection of the primary tumour can limit post-surgical disease progression and improve survival.

While these findings establish the therapeutic potential of endothelin receptor blockade in combination with T cell-based cellular immunotherapy, several aspects warrant further investigation. Although therapeutic benefit was demonstrated across two tumour models and two distinct cellular immunotherapy platforms, further validation in additional tumour settings will be important to define the broader applicability of this therapeutic strategy. Moreover, although this study identifies biological processes and intratumoral T cell states associated with improved therapeutic efficacy, the mechanisms through which ERB enhances adoptively transferred T cell responses remain to be fully elucidated. The high-dimensional phenotypic profiling was performed at a single experimental endpoint and therefore identifies associations rather than causal relationships. Longitudinal studies will be important to define how the tumour-control-associated T cell populations emerge and evolve during treatment and to determine their functional contribution to therapeutic efficacy and durable tumour control. Tracking transferred T cells independently from endogenous populations will additionally be important for determining how ERB influences their tumour accumulation and persistence, and whether transferred T cells contribute to the intratumoral T cell states associated with tumour control identified here. Finally, although resistance to tumour rechallenge is consistent with the development of durable antitumour immune memory, the immune cell populations and mechanisms responsible for this protection remain to be determined.

In conclusion, our findings identify the endothelin receptor pathway as a promising therapeutic target for enhancing cellular immunotherapy and provide new insights into the intratumoral T cell states associated with successful antitumour immunity. By increasing tumour control responses and, in combination with ICI, promoting durable tumour control and long-term protection against tumour rechallenge, this therapeutic strategy addresses two major challenges facing cellular immunotherapy. Importantly, endothelin receptor antagonists are clinically approved for non-oncological indications, while immune checkpoint inhibitors and cellular immunotherapies are established cancer therapies, potentially lowering some of the translational barriers to evaluating these combinations in cancer patients. Together, these findings provide a strong rationale for the clinical investigation of endothelin receptor blockade as a strategy to improve the efficacy and durability of T cell-based cancer immunotherapies across distinct therapeutic platforms and tumour settings.

## MATERIALS AND METHODS

### Cells

4T1 (ATCC CRL-2539) murine mammary carcinoma cells, A20 murine B cell lymphoma cells (ATCC TIB-208) were purchased from ATCC and RENCA murine renal adenocarcinoma cells were kindly provided by Andreas Bikfalvi (Angiogenesis and Cancer Microenvironment Laboratory, University of Bordeaux, France). All cell lines were routinely confirmed to be free of Mycoplasma contamination. Raji B cells were kindly provided by the Antibody and Vaccine group, University of Southampton. 4T1, RENCA, A20, Raji B cells were maintained in RPMI-1640 (Gibco) supplemented with 10% FBS (Gibco), penicillin (100 U/mL; Gibco), streptomycin sulfate (100 μg/mL; Gibco), L-glutamine (2 mM; Gibco), sodium pyruvate (1 mM; Gibco) and 1× non-essential amino acids (NEAA; Gibco). Primary T cells were maintained in the same complete RPMI-1640 medium additionally supplemented with 55 μM 2-mercaptoethanol (Gibco, Cat. #21985-023).

### Flow cytometry

For all flow cytometry experiments, cell suspensions were stained with fixable eFluor 780 viability dye (Invitrogen), followed by surface antibody staining. The following antibodies against murine molecules were used: CD45-PerCP (Cat. #103130), CD3-Spark Blue 550 (Cat. #100260), CD4-Brilliant Violet 750 (Cat. #100467), CD8-Brilliant Violet 570 (Cat. #100740), CD62L-PE/Dazzle 594 (Cat. #104448), CD44-Brilliant Violet 570 (Cat. #103057), CD19-Alexa Fluor 488 (Cat. #115521), CD49b-PE/Cyanine 7 (Cat. #108922), CD2-PE (Cat. #100108), CD28-Brilliant Violet 421 (Cat. #102127), CD279/PD-1-Alexa Fluor 647 (Cat. #135230), CD69-PE/Dazzle 594 (Cat. #104536), CD25-PE/Cyanine 7 (Cat. #101916), TIM-3-PE/Fire 640 (Cat. #119749), CD223/LAG-3-Brilliant Violet 785 (Cat. #125219), CD45-Brilliant Violet 510 (Cat. #103138), CD3-FITC (Cat. #100204), CD4-Brilliant Violet 570 (Cat. #100542), CD8α-Spark Blue 550 (Cat. #100780), CD25-Brilliant Violet 750 (Cat. #102077), CD127-Brilliant Violet 711 (Cat. #135035), CD49b-PerCP/Cyanine 5.5 (Cat. #108916), CD62L-Brilliant Violet 785 (Cat. #104440), CD44-Brilliant Violet 605 (Cat. #103047), CD45R/B220-PerCP (Cat. #103234), CD28-PE/Cyanine 5 (Cat. #102108), 4-1BB/CD137-PE/Cyanine 7 (eBioscience, Cat. #MA546776), CD39-PE/Dazzle 594 (Cat. #143811), CD223/LAG-3-Brilliant Violet 650 (Cat. #125227) and CD279/PD-1-Alexa Fluor 647 (Cat. #135229). All antibodies were purchased from BioLegend unless otherwise stated.

### In vitro activation and proliferation monitoring of splenocyte-and LN-derived murine T cells

Murine Pan T cells were isolated from splenocytes or tumour-draining lymph nodes by magnetic separation according to the manufacturer’s protocol (Miltenyi Biotec, Cat. #130-095-130). For proliferation assessment, T cells were labelled with CFSE and stimulated with plate-bound anti-CD3 alone or in combination with plate-bound anti-CD2 antibodies, with unstimulated cells maintained as resting controls. Activation and proliferation were assessed by flow cytometric analysis of CD69 and CD25 expression and CFSE dilution, respectively. Following stimulation, activated T cells were expanded in complete RPMI medium supplemented with IL-2, phenotyped by flow cytometry and used for *in vitro* functional assays or adoptive T cell transfer as indicated.

### Generation of adoptively transferred T cells

Female BALB/c donor mice aged 6-8 weeks were orthotopically implanted with 5 × 10⁴ 4T1 cells into the third mammary fat pad. At 10-13 days after tumour implantation, tumour-draining inguinal lymph nodes (TDLNs) were harvested and processed into single-cell suspensions. Pan T cells were isolated from freshly harvested TDLNs by immunomagnetic separation (Miltenyi Biotec, Cat. #130-095-130). Isolated T cells were activated ex vivo using plate-bound anti-CD3 and anti-CD2 antibodies and subsequently expanded in IL-2-containing culture medium. T-cell activation was assessed by flow cytometric analysis of CD69 and CD25 expression. Expanded T cells were phenotyped by flow cytometry to determine the proportions of CD4⁺ and CD8⁺ T cells.

### Syngeneic 4T1 Tumour Model and Treatment Protocols

The orthotopic 4T1 mammary tumour model was established by implanting 5 × 10⁴ 4T1 cells into the third mammary fat pad of female BALB/c mice aged 6-8 weeks. Mice were obtained from The Cyprus Institute of Neurology and Genetics. All *in vivo* experiments were conducted in accordance with animal welfare regulations and guidelines of the Republic of Cyprus and the European Union (European Directive 2010/63/EU and Cyprus Legislation for the Protection and Welfare of Animals, Laws 1994-2013). Experiments were conducted under project licence #CY/EXP/PR.L04/2023 and personal licence #CY/EXP/P.L07/2022 issued by the Cyprus Veterinary Services.

For the *in vivo* 4T1 ACT studies, tumour-bearing recipient mice received a single intravenous infusion of 1 × 10⁶ viable T cells when mean tumour volume was approximately 100 mm³. Mice were treated with ACT alone or in combination with bosentan-mediated endothelin receptor blockade (ERB) (Selleckchem, Cat. #S3051) and/or immune checkpoint inhibition (ICI) using anti-PD-1 (Bio X Cell, Cat. #BP0146) and anti-CTLA-4 (Bio X Cell, Cat. #BP0164) antibodies. Bosentan was administered intraperitoneally at 1 mg/kg once daily in a vehicle comprising 2% Dimethyl sulfoxide (DMSO), 30% PEG300, 2% Tween-80 and 66% water. Corresponding control mice received an equivalent volume of vehicle according to the same administration schedule. The 1 mg/kg bosentan dose was selected based on dose-optimisation studies demonstrating favourable effects on tumour stiffness, vascular function and oxygenation in both 4T1 and E0771 triple-negative breast cancer models^42^. ERB was initiated before T-cell transfer and, following omission on the day of ACT, was resumed the following day and continued once daily thereafter, according to the treatment schedule (Fig. S3). ICI was administered intraperitoneally as a combination of anti-PD-1 (10 mg/kg) and anti-CTLA-4 (5 mg/kg)^42,64,65^, beginning one day after ACT, with two additional doses administered at three-day intervals. Corresponding single-treatment groups received the respective agents according to the same schedules. Tumour growth was monitored longitudinally using digital callipers, and tumour volume was calculated as V = 4/3 × π × (d/2)³, where d represents the mean of the two measured tumour dimensions. Mouse body weight was monitored longitudinally throughout the experimental period. Mice were euthanised when tumour volume reached or exceeded the predefined humane endpoint of 1200 mm³ at scheduled tumour assessment or if body weight decreased by ≥20% relative to starting body weight. Lung metastases were assessed at the experimental or humane endpoint, as indicated. For long-term survival studies, mice were monitored for up to 110 days following primary tumour inoculation. Long-term survivors were subsequently subjected to tumour rechallenge as described below.

### Post-surgical 4T1 tumour model and treatment

The orthotopic 4T1 tumour model was established and TDLN-derived T cells for ACT were generated as described above. Briefly, mice were implanted with 5 × 10⁴ 4T1 cells into the third mammary fat pad, and ACT products were generated from TDLN-derived T cells following *ex vivo* anti-CD3/anti-CD2 activation and expansion. On day 14 following tumour inoculation, established primary tumours were surgically resected. Following tumour resection, mice were assigned to either a surgery-only control group or post-operative combination therapy comprising ACT, bosentan-mediated endothelin receptor blockade (ERB) and immune checkpoint inhibition (ICI), administered as described above starting on Day 17 with ERB. Mice were subsequently monitored for survival until the predefined experimental or humane endpoints were reached.

### Tumour response classification

Individual tumour responses were classified according to tumour growth kinetics during treatment and longitudinal follow-up, rather than tumour status at a single fixed timepoint. These study-defined criteria were developed to capture treatment-associated tumour control and response durability in the experimental models and were not intended to represent clinical RECIST criteria. To establish a response threshold, the lowest tumour volume observed among control mice at the end of the treatment period was used as the reference threshold. Complete tumour regression was defined as no measurable tumour (tumour volume = 0) for at least two consecutive monitoring timepoints.

Progressive disease (PD) was defined by an overall pattern of continued tumour growth throughout the experimental observation period. Stable disease (SD) was defined by relatively stable tumour volume, with ≤20% variation between consecutive measurements for at least three consecutive timepoints, without subsequent complete tumour regression. Partial tumour control (PTC) was defined by tumour volume falling below the response threshold at the end of the treatment period without complete tumour regression, followed by sustained tumour progression during longitudinal follow-up. Complete response (CR) was defined as complete tumour regression followed by tumour recurrence within the experimental observation period, whereas maintained complete response (MCR) was defined as complete tumour regression maintained without recurrence throughout the remainder of the experimental observation period. PTC, CR and MCR were collectively considered tumour control responses, reflecting treatment-associated tumour control ranging from reduced tumour progression to complete and maintained tumour regression.

### Cytotoxicity assays

The cytotoxic activity of expanded 4T1 TDLN-derived T cells against 4T1 tumour cells was assessed using the CyQUANT™ LDH Cytotoxicity Assay Kit according to the manufacturer’s instructions (Invitrogen, Cat. #C20300). The murine renal adenocarcinoma cell line RENCA was included as an irrelevant target-cell control. Target cells (5 × 10⁴) were co-cultured with expanded 4T1 TDLN-derived T cells for 48 h at effector-to-target (E:T) ratios of 1:1 or 5:1. The cytotoxic activity of anti-CD19 CAR T cells was assessed using the same LDH-based assay. Anti-CD19 CAR T cells were co-cultured with mouse A20 B-cell lymphoma cells or antigen-mismatched human Raji B cells (5 × 10⁴) at E:T ratios of 1:1 or 5:1 for 48 h. Four independent CAR T-cell cytotoxicity experiments were performed. LDH release was measured in culture supernatants, with target cells alone used to determine spontaneous LDH release and lysed target cells used to determine maximum LDH release. Percentage cytotoxicity was calculated as [(experimental release - spontaneous release)/(maximum release - spontaneous release)] × 100. Absorbance was measured at 490 nm with a 590-nm reference wavelength.

### Generation of single-cell suspensions from harvested 4T1 tumours for flow cytometry

Harvested 4T1 tumours were weighed, mechanically minced and enzymatically digested with Liberase TL (Sigma-Aldrich, Cat. #05401020001) supplemented with DNase I for 15-20 min at 37°C with agitation. Digestion was stopped by addition of PBS or culture medium supplemented with 10% Fetal Bovine Serum (FBS). Tumour suspensions were passed through a sterile cell strainer, with remaining tissue mechanically dissociated to maximise cell recovery. The resulting cell suspensions were centrifuged, resuspended in PBS, counted and processed for flow cytometric staining.

### High-dimensional phenotypic analysis

Flow cytometry standard (FCS) files were analysed using FlowJo software (v10.8.1). Following exclusion of debris based on forward-and side-scatter properties, singlets were identified using FSC-A versus FSC-H and SSC-A versus SSC-H gating, and dead cells were excluded using Fixable Viability Dye eFluor 780. Subsequent analyses were restricted to CD45⁺CD49b⁻B220⁻CD3⁺ tumour-infiltrating T cells. High-dimensional analysis was performed using t-distributed stochastic neighbour embedding (tSNE) followed by Phenograph clustering. tSNE was performed in FlowJo using the optSNE implementation with 1,000 iterations and a perplexity of 30, using the exact (vantage-point tree) nearest-neighbour and Barnes-Hut gradient algorithms. Phenograph clustering was performed using expression of the following markers: PD-1, CD4, CD44, LAG-3, CD127, CD25, CD62L, CD2, CD28, 4-1BB, CD39 and CD8α. Clusters were manually annotated based on lineage, activation, memory and co-stimulatory marker-expression patterns visualised using heatmaps and tSNE projections. Frequencies of individual clusters were calculated for each sample and correlated with tumour volume measured on the day of tumour harvest, one day after completion of all therapeutic regimens, to identify T-cell populations positively or negatively associated with tumour burden.

### Immunofluorescence staining and image analysis

Immunofluorescence staining and image analysis were performed as previously described^42^. In brief, tumours were excised, washed twice in 1× PBS for 10 min and fixed in 4% paraformaldehyde (PFA) for 24 h at 4°C. Following fixation, samples were washed twice in 1× PBS for 10 min, dehydrated through successive steps of ethanol and xylene, and embedded in paraffin. Tissues were then sectioned at 7 μm using a rotary microtome (Accu-Cut SRM 200; Sakura). Sections were mounted onto microscope slides, dried overnight at 37°C, deparaffinized and rehydrated before immunofluorescence staining.

For analysis of tumour-infiltrating T-cell populations, sections were incubated overnight at 4°C with either rabbit anti-CD4 (clone EPR19514, Abcam, 1:200) together with rat anti-CD3 (clone 17A2, BioLegend, 1:100) or rat anti-CD8α (clone 4SM15, Thermo Fisher Scientific/eBioscience, 1:100) together with rabbit anti-granzyme B (clone EPR22645-206, Abcam, 1:100). Following washing, sections were incubated for 2 h at room temperature with species-appropriate Alexa Fluor 488- or Alexa Fluor 647-conjugated secondary antibodies (Invitrogen, 1:400). Nuclei were counterstained with DAPI before mounting with ProLong Gold Antifade Mountant (Invitrogen).

Fluorescence images were acquired using a Leica Stellaris 5 confocal microscope with identical acquisition settings for all samples within each staining panel. For histological analyses, 2-3 tumours per experimental group were selected based on tumour size at the time of harvest, to represent the average tumour size within the respective experimental group. Five fields of view were analysed per tumour. Image analysis was performed in MATLAB (MathWorks, Natick, MA, USA) using a combination of custom-developed and built-in algorithms as previously described^42^. Identical fluorescence intensity thresholds were applied to all images for each fluorescence channel to minimise background signal and ensure consistent analysis across experimental groups. Quantification was performed using area fraction rather than cell counting. Specifically, CD8 abundance was calculated as the fraction of CD8-positive pixels relative to DAPI-positive pixels, while CD4⁺CD3⁺ and CD8⁺granzyme B⁺ populations were quantified as the fraction of co-localised double-positive pixels relative to DAPI-positive pixels. Co-localisation was determined based on pixel overlap between the corresponding fluorescence channels.

### Anti-CD19 CAR T cell generation

The anti-mouse CD19 CAR was a second-generation CAR comprising an anti-mouse CD19 scFv (clone 1D3), a murine CD8 stalk, a murine CD28 costimulatory domain and a murine CD3ζ signalling domain. The mock CAR comprised an anti-human CD19 scFv (clone FMC63), a human CD8 stalk and transmembrane domain, a human 4-1BB costimulatory domain and a human CD3ζ signalling domain. Both CAR constructs were encoded in SFG retroviral vectors together with a Thy1.1 marker in a Thy1.1-2A-CAR configuration.

Retroviral vectors encoding the anti-mouse CD19 and mock CAR constructs were produced using Phoenix Eco packaging cells transfected with the corresponding envelope and transfer plasmids using GeneJuice transfection reagent (Merck). Viral supernatants were harvested following transfection, clarified and used for subsequent T-cell transduction. Murine T cells were isolated from splenocytes and activated *ex vivo* using plate-bound anti-CD3 and anti-CD2 antibodies as described above. Activated T cells were transduced with the corresponding retroviral vectors using Retronectin-assisted transduction and subsequently expanded in IL-2-containing culture medium. Prior to *in vivo* use, CAR expression and the composition of the resulting T-cell populations were assessed by flow cytometry.

### A20 B cell lymphoma mouse model

The ectopic A20 B-cell lymphoma model was established by subcutaneous implantation of 0.4 × 10⁶ A20 cells in the region of the third mammary fat pad of female BALB/c mice aged 6–8 weeks. For the *in vivo* CAR T-cell studies, tumour-bearing recipient mice received a single intravenous infusion of 1 × 10⁶ viable T cells. Mice received either Mock CAR T cells or anti-CD19 CAR T cells, with anti-CD19 CAR T-cell therapy administered alone or in combination with bosentan-mediated endothelin receptor blockade (ERB) and/or immune checkpoint inhibition (ICI), using the same agents and formulations described above for the 4T1 ACT studies (bosentan, 1 mg/kg; anti-PD-1, 10 mg/kg; anti-CTLA-4, 5 mg/kg). ERB was initiated when tumours reached an average diameter of approximately 5 mm. Following omission on the day of CAR T-cell transfer, ERB was resumed the following day and continued once daily thereafter. ICI was initiated one day after CAR T-cell transfer and administered at three-day intervals, according to the treatment schedule detailed in Fig. S6. Tumour growth was monitored longitudinally using digital callipers, and tumour volume was calculated as V = 4/3 × π × (d/2)³, where d represents the mean of the two measured tumour dimensions. Mice were euthanised when tumour volume reached or exceeded the predefined humane endpoint of 1200 mm³ at scheduled tumour assessment or if body weight decreased by ≥20% relative to starting body weight.

### 4T1 and A20 tumour rechallenge in vivo assay

Tumour rechallenge experiments were performed to assess resistance to tumour re-establishment in mice that achieved a maintained complete response (MCR) following triple-combination treatment comprising T-cell therapy, endothelin receptor blockade (ERB) and immune checkpoint inhibition (ICI).

In the 4T1 model, MCR mice were rechallenged approximately 110 days after primary tumour inoculation with 5 × 10⁴ viable 4T1 cells implanted orthotopically into the contralateral mammary fat pad. In the A20 model, MCR mice were similarly rechallenged approximately 110 days after primary tumour inoculation with 0.4 × 10⁶ viable A20 cells by ectopic subcutaneous implantation in the contralateral mammary region. Tumour-naïve, untreated, age- and sex-matched mice were challenged in parallel with the corresponding tumour cells as controls. No additional therapeutic treatment was administered following rechallenge.

Mice were monitored longitudinally for tumour development, tumour volume, body weight, clinical condition and survival until the predefined experimental endpoint or humane endpoint criteria were reached. Tumour dimensions were measured using digital callipers, and tumour volume was calculated as V = 4/3 × π × (d/2)³, where d represents the mean of the two measured tumour dimensions.

### Statistical analyses

All statistical analyses were performed using GraphPad Prism software (version 10.4.1). Unless otherwise specified in the figure legends, comparisons between two groups were performed using two-tailed unpaired *t*-tests with Welch’s correction. Associations between T-cell cluster frequencies and tumour volume were assessed using two-tailed Pearson correlation analysis. Survival curves were estimated using the Kaplan-Meier method and compared using the log-rank (Mantel-Cox) test. Statistical significance was defined as *P* < 0.05.

## ACKNOWLEDGEMENTS

The authors thank Malika Hoekx and Foongjun Yap for their assistance and support during murine CAR T-cell training. The authors thank the staff of the CING mouse facility for their assistance with animal husbandry and support of the *in vivo* studies. The authors also thank Dr Chryso Pieridou and the Human Resources team at CSHM for their support. This project has received funding from the Cyprus Cancer Research Institute (CCRI) under Funding Agreement No. CRI_2020_FA_LE_101 through the BRIDGES competitive funding call, with additional support from the Karaiskakio Foundation and The Andreas and Shirley Kramvis Foundation.

## Author contributions

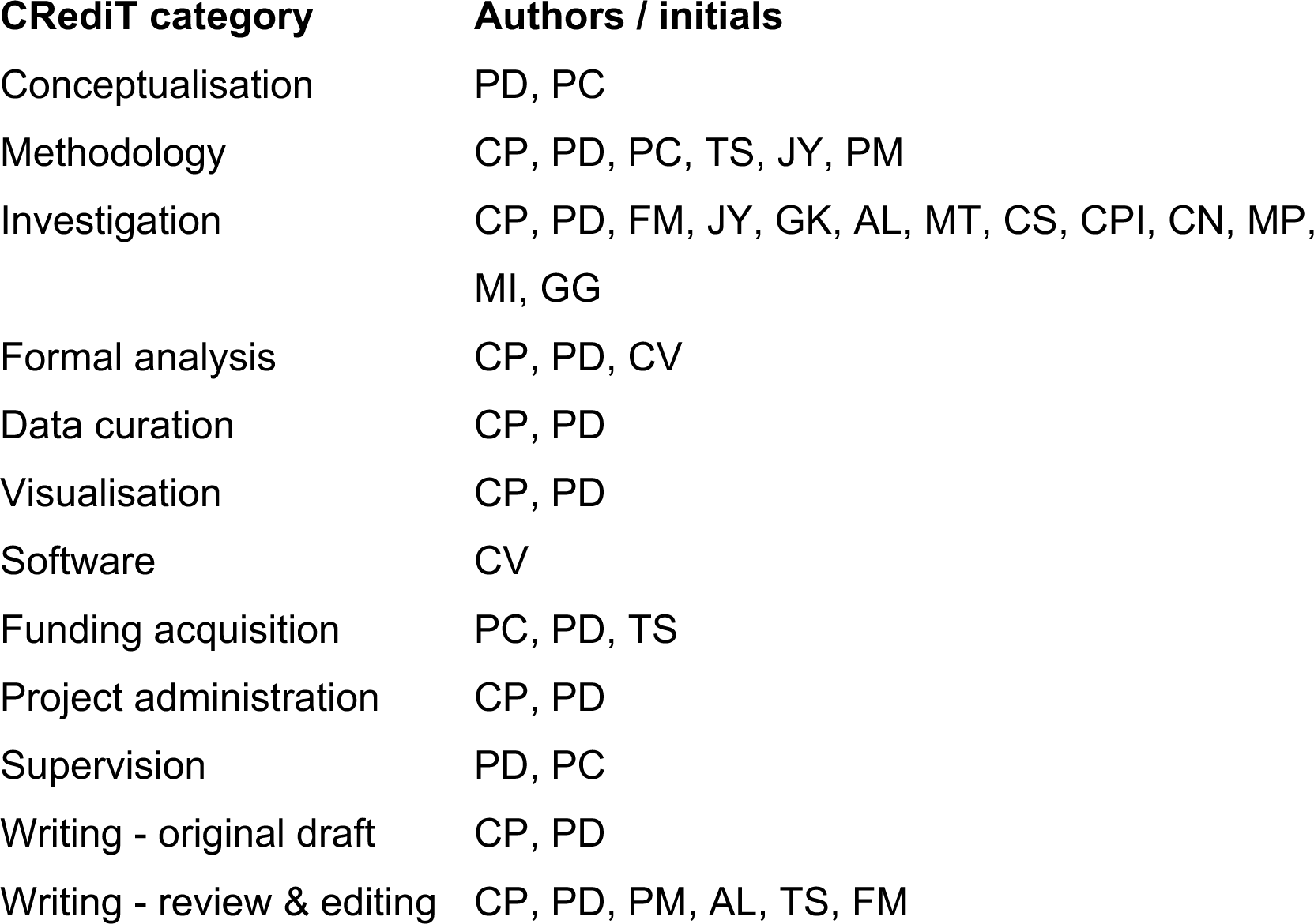

## Declaration of Generative AI and AI-Assisted Technologies

During the preparation of this manuscript, the authors used ChatGPT (OpenAI) to assist with language editing and to improve the clarity and readability of author-generated text. Following the use of this tool, the authors reviewed and revised the content as appropriate and take full responsibility for the content of the published article.

