## Supplementary Information for "Endothelin receptor blockade potentiates adoptive T cell and CAR T cell therapies"

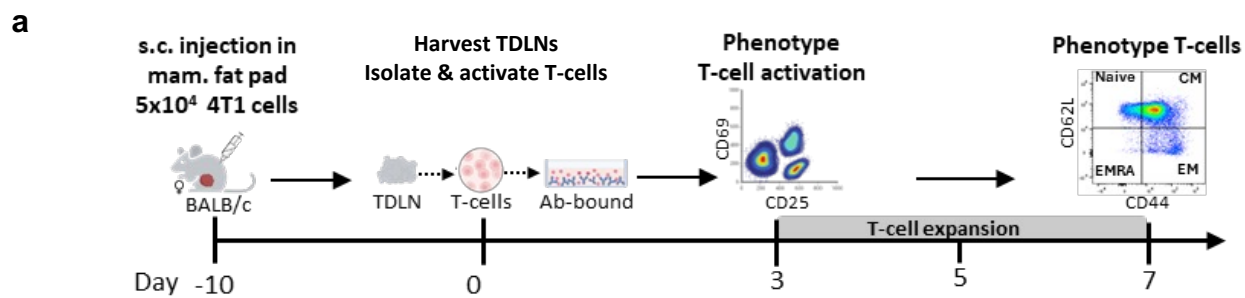

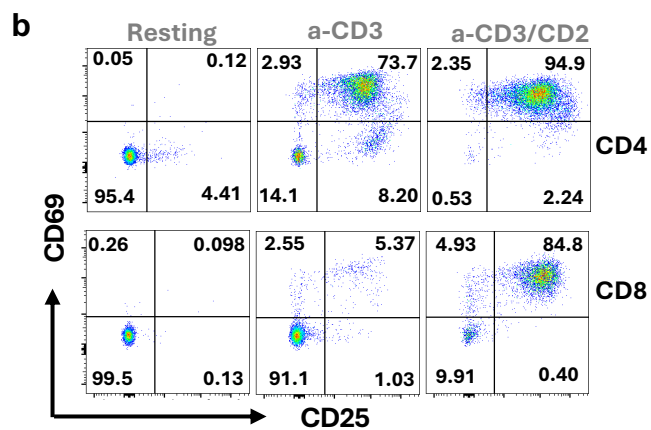

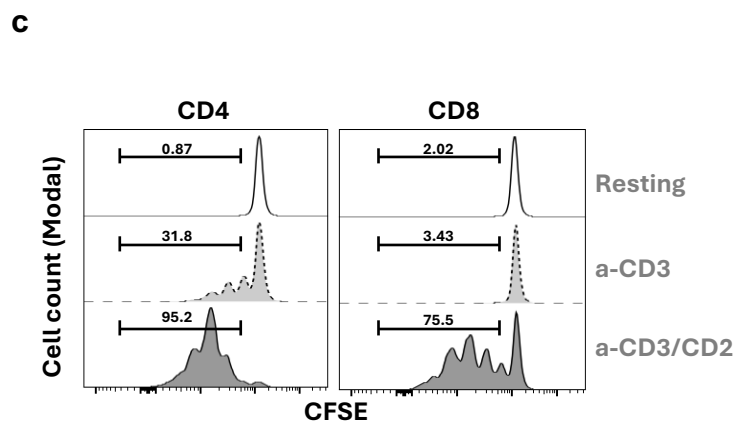

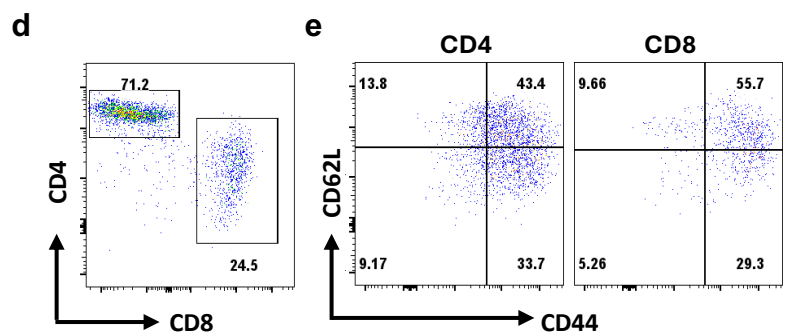

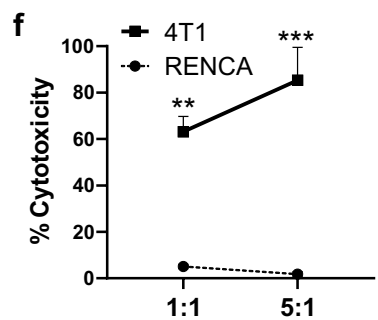

**Fig. S1: Generation and functional characterisation of the 4T1 tumour-reactive T cell product used for adoptive cell therapy.**

**(a)** Schematic representation of the generation and characterisation of the tumour-draining lymph node (TDLN)-derived T-cell product used for adoptive cell therapy in the 4T1 model.

**(b)** Representative flow cytometry plots showing T-cell activation based on CD69 and CD25 expression in resting and stimulated 4T1 TDLN-derived pan T cells at 72 h post-stimulation.

**(c)** Proliferation of CFSE-labelled pan T cells following 72 h of stimulation with anti-CD3 or anti-CD3/CD2. CFSE profiles are shown separately for CD4<sup>+</sup>CD3<sup>+</sup> and CD8<sup>+</sup>CD3<sup>+</sup> T cells following gating on live singlets.

**(d)** Representative flow cytometry plots showing the CD4:CD8 T cell composition of the culture at Day 7 post-stimulation.

**(e)** Naïve/memory phenotype of CD4<sup>+</sup> and CD8<sup>+</sup> T-cell subsets at Day 7, based on CD62L and CD44 expression.

**(f)** Cytotoxic activity of the Day 7 T-cell product following 48-h co-culture with 4T1 or RENCA target cells at effector-to-target (E:T) ratios of 1:1 and 5:1. RENCA cells were included as an irrelevant tumour target control.

**a**

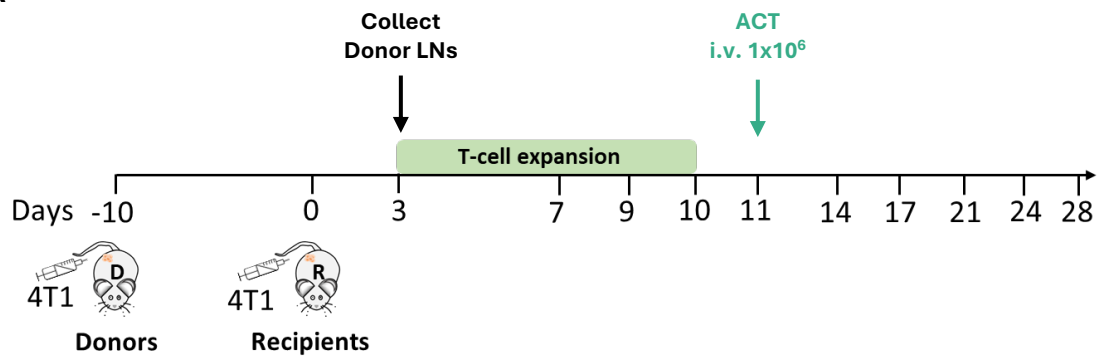

**b**

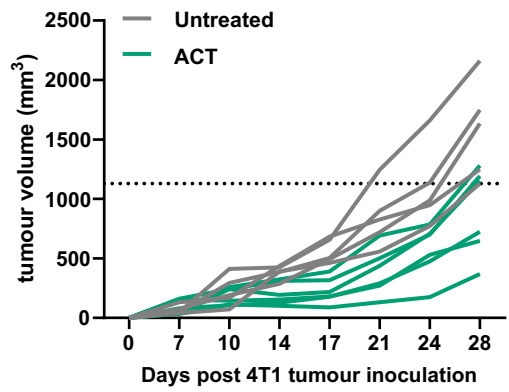

**c**

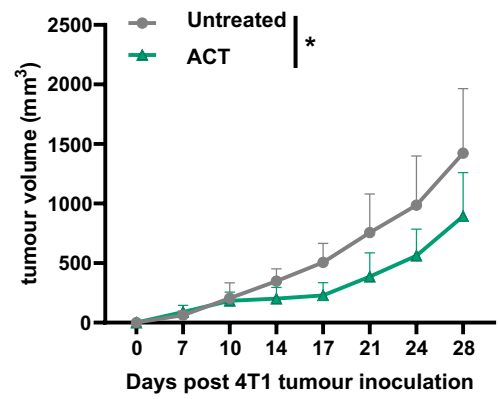

**d**

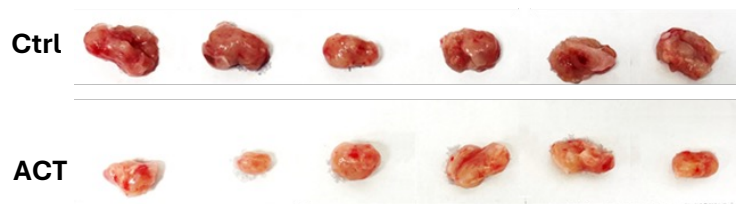

**e**

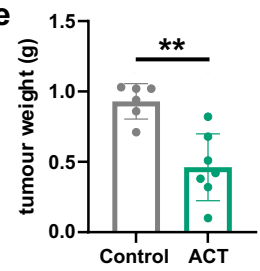

**Fig. S2: Therapeutic activity of 4T1 tumour-draining lymph node-derived T cells following adoptive transfer.**

**(a)** Schematic representation of the 4T1 adoptive T-cell therapy model. 4T1 cells were implanted into the mammary fat pad of 6-8-week-old female BALB/c mice. In vitro-expanded 4T1 tumour-draining lymph node (TDLN)-derived T cells were administered intravenously when tumours reached an average volume of approximately 100 mm<sup>3</sup>.

**(b-c)** Individual **(b)** and mean **(c)** tumour growth curves of 4T1 tumour-bearing mice receiving no treatment or adoptive T-cell therapy (ACT).

**(d)** Representative images of tumours collected at Day 28, when the first mice reached a tumour volume of approximately 1,200 mm<sup>3</sup>.

**(e)** Tumour weights at Day 28.

Statistical significance was determined using a two-tailed unpaired t-test with Welch's correction. \*$P < 0.05$; \*\*$P < 0.01$.

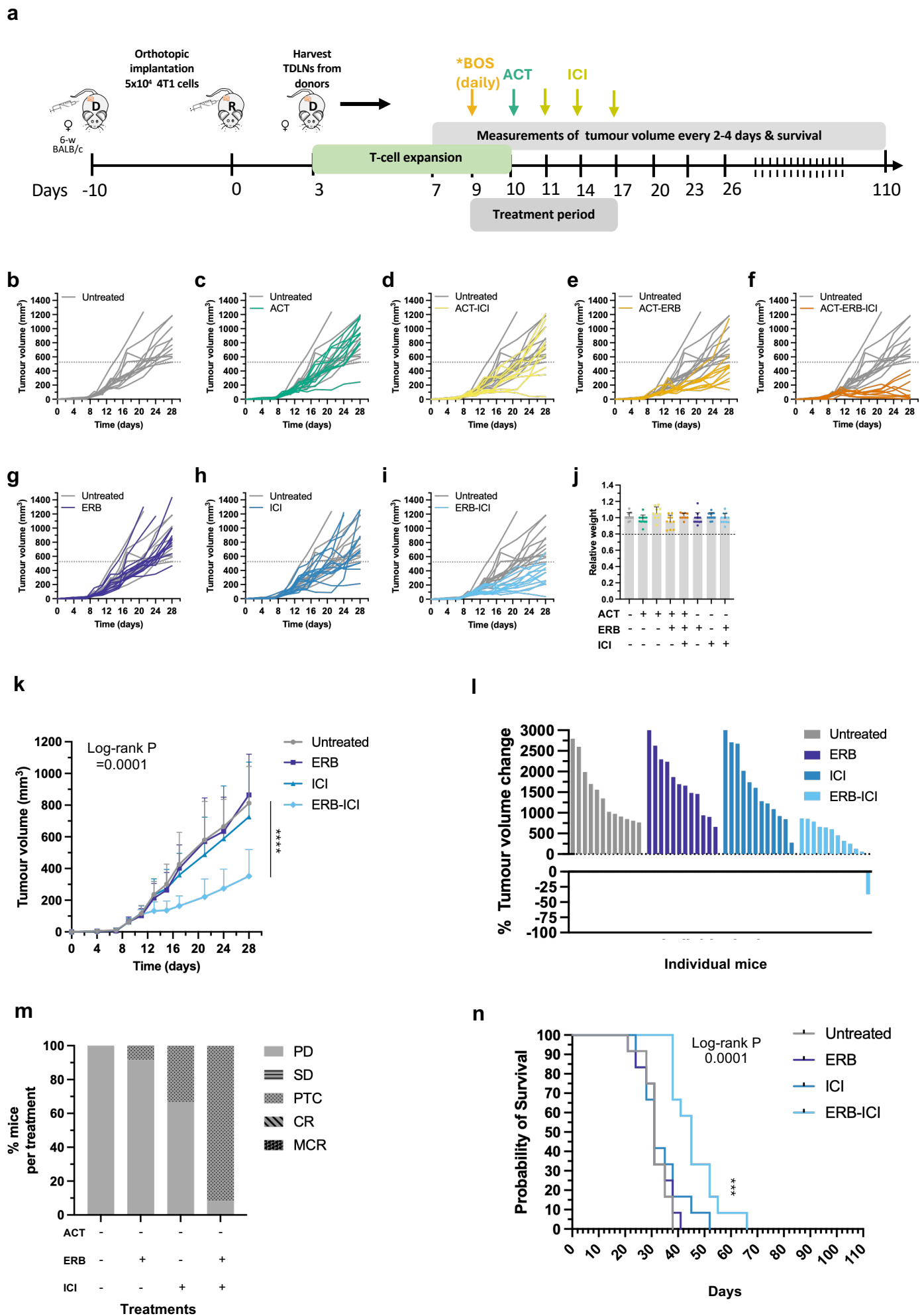

**Fig. S3: Endothelin receptor blockade enhances adoptive T cell therapy alone and in combination with immune checkpoint inhibition.**

**(a)** Experimental schema showing orthotopic implantation of 4T1 tumour cells, generation and expansion of tumour-draining lymph node-derived T cells, and treatment schedule for ACT, bosentan-mediated endothelin receptor blockade (ERB), and immune checkpoint inhibition (ICI). ERB treatment was initiated once tumours reached an average diameter of approximately 5 mm, which typically occurred around Day 9 after tumour inoculation. Bosentan was administered daily for 9 days, with the exception of the day of ACT administration. ACT was administered one day after initiation of ERB, and ICI was administered one day after ACT followed by additional doses at 3-day intervals. Tumour growth and survival were monitored until study completion.

**(b-i)** Individual tumour growth curves for mice receiving Control, ACT, ACT-ICI, ACT-ERB, ACT-ERB-ICI, ERB, ICI, or ERB-ICI treatment. Each line represents an individual mouse. Dashed lines indicate the humane endpoint tumour volume.

**(j)** Relative body weight following completion of treatment, calculated for each mouse with paired measurements as body weight at Day 17 divided by body weight at Day 9. Individual mice and mean ± SD are shown. The dashed line at 0.8 indicates the predefined humane endpoint corresponding to a 20% reduction from starting body weight.

**(k)** Mean tumour growth kinetics in ERB alone, ICI alone, or ERB-ICI treatment groups compared with untreated controls.

**(l)** Waterfall plot showing the percentage change in tumour volume for individual mice in ERB alone, ICI alone, or ERB-ICI treatment groups compared with untreated controls.

**(m)** Tumour response classification. Individual responses were classified using study-defined criteria based on tumour growth kinetics during treatment and longitudinal follow-up. Responses were classified as progressive disease (PD), stable disease (SD), partial tumour control (PTC), complete response (CR) or maintained complete response (MCR). Data are presented as the percentage of mice in each response category. Tumour control responses comprise PTC, CR and MCR. See Methods for the definition and evaluation of each response category.

**(n)** Kaplan-Meier survival analysis of mice in ERB alone, ICI alone, or ERB-ICI treatment groups compared with untreated controls.

### Re-challenge study

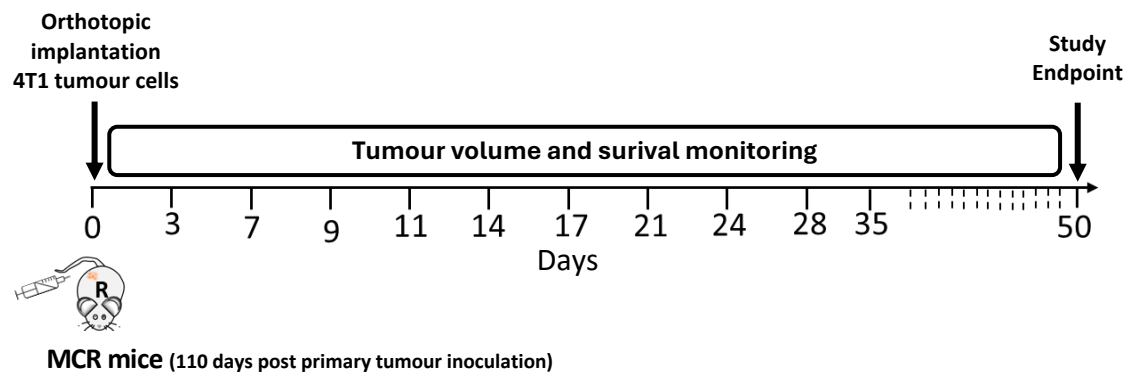

**Fig. S4: Experimental schema of the tumour rechallenge study.**

Mice achieving maintained complete responses (MCRs) following ACT-ERB-ICI treatment in the primary 4T1 tumour experiment were rechallenged with the same number of 4T1 tumour cells 110 days after primary tumour inoculation. Tumour cells were implanted orthotopically into the contralateral mammary fat pad. Tumour growth and survival were monitored for 50 days following rechallenge. Treatment-naïve mice challenged in parallel served as controls.

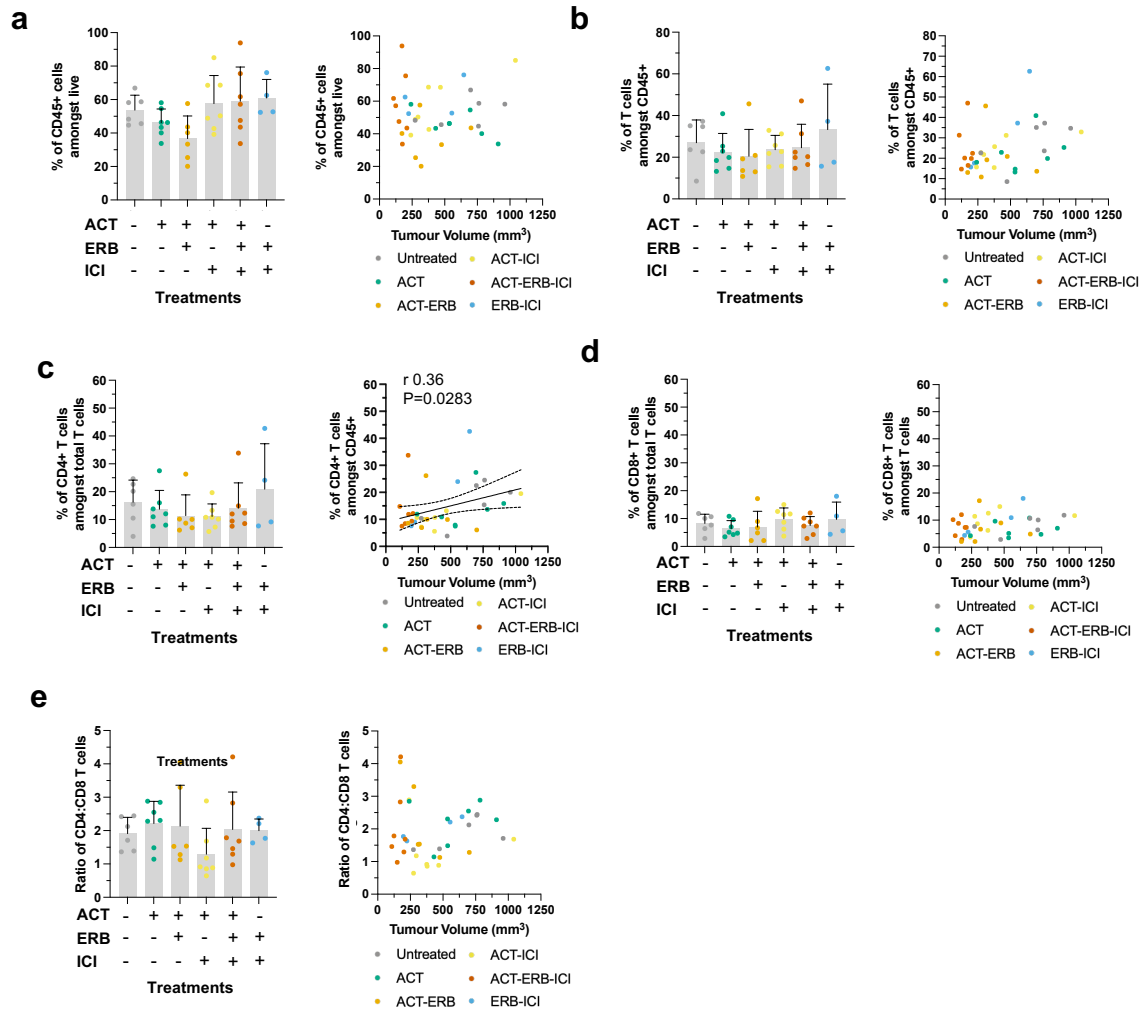

**Fig. S5: Conventional flow cytometric analyses of tumour-infiltrating immune populations and their relationship with tumour burden.**

- **(a)** Frequency of CD45<sup>+</sup> immune cells within tumours across treatment groups (left) and correlation between CD45<sup>+</sup> cell frequency and tumour volume at harvest (right).
- **(b)** Frequency of total CD45<sup>+</sup>CD3<sup>+</sup> T cells within tumours across treatment groups (left) and correlation between CD45<sup>+</sup>CD3<sup>+</sup> T cell frequency and tumour volume at harvest (right).
- **(c)** Frequency of CD4<sup>+</sup> T cells within tumours across treatment groups (left) and correlation between CD4<sup>+</sup> T cell frequency and tumour volume at harvest (right). A significant positive correlation between tumour volume and total CD4<sup>+</sup> T-cell frequency was observed ($r = 0.36$, $R^2 = 0.1301$, $P = 0.0283$).
- **(d)** Frequency of CD8<sup>+</sup> T cells among total tumour-infiltrating T cells across treatment groups (left) and correlation between CD8<sup>+</sup> T cell frequency and tumour volume at harvest (right).
- **(e)** Ratio of CD4:CD8 T cells within tumours across treatment groups (left) and correlation between the CD4:CD8 ratio and tumour volume at harvest (right).

For treatment-group comparisons, bars represent mean ± SD and each symbol represents an individual mouse. Associations with tumour volume were assessed using Pearson correlation analysis. Tumour volumes correspond to measurements obtained one day after completion of all treatments.

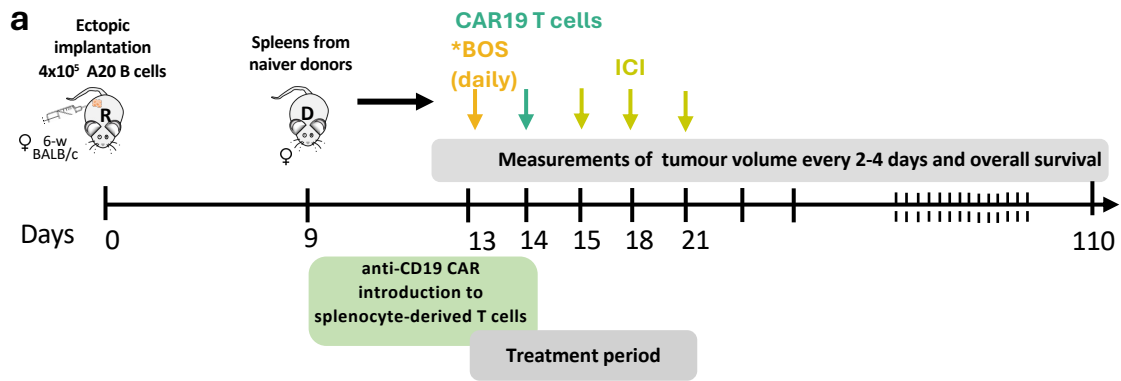

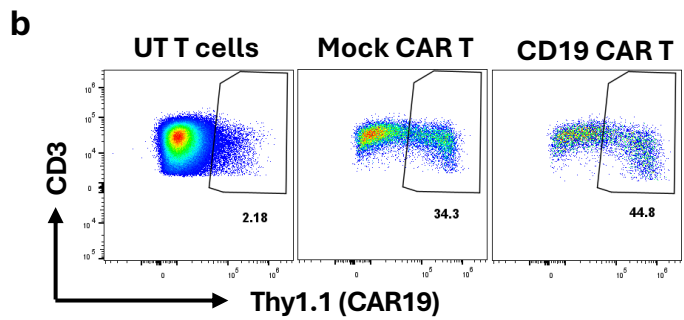

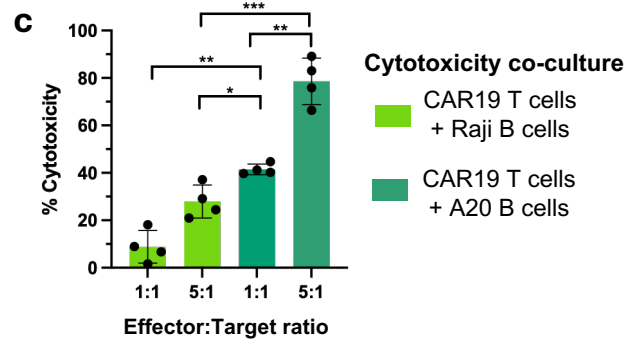

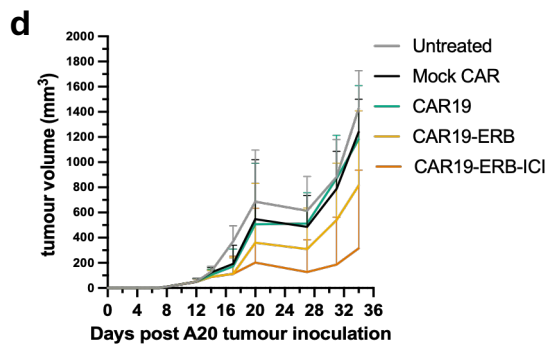

**Fig. S6: Experimental design, functional characterisation and longitudinal tumour growth analysis of anti-CD19 CAR T-cell therapy.**

**(a)** Experimental schematic showing A20 tumour implantation, generation and adoptive transfer of anti-CD19 CAR T cells (CAR19), bosentan-mediated endothelin receptor blockade (ERB), immune checkpoint inhibition (ICI), tumour monitoring and study endpoints. Treatment timing and administration schedules are indicated in the experimental schema.

**(b)** Representative flow cytometric analysis of Thy1.1 expression as a marker of CAR T-cell transduction prior to adoptive transfer, with approximately 50% of T cells expressing the Thy1.1 transduction marker.

**(c)** *In vitro* cytotoxicity of anti-CD19 CAR T cells against human Raji B cells and mouse A20 B-cell lymphoma cells at effector-to-target (E:T) ratios of 1:1 and 5:1, assessed using an LDH release assay. Each data point represents an independent CAR19 T cell cytotoxicity experiment ($n = 4$ independent experiments). CAR19 T cells exhibited significantly greater cytotoxicity against A20 cells than against the antigen-mismatched Raji cells at both 1:1 ($P = 0.0013$) and 5:1 ($P = 0.0003$) E:T ratios. Cytotoxicity against A20 cells was also significantly increased at an E:T ratio of 5:1 compared with 1:1 ($P = 0.0036$). Statistical comparisons were performed using two-tailed unpaired *t*-tests with Welch's correction.

**(d)** Mean tumour growth curves corresponding to the individual tumour growth trajectories shown in Fig. 5a. Mice received mock CAR T cells, anti-CD19 CAR T cells alone (CAR19), CAR19-ERB, or CAR19-ERB-ICI. Tumour volumes were measured every 3–4 days. Data are presented as mean tumour volume ± SD.

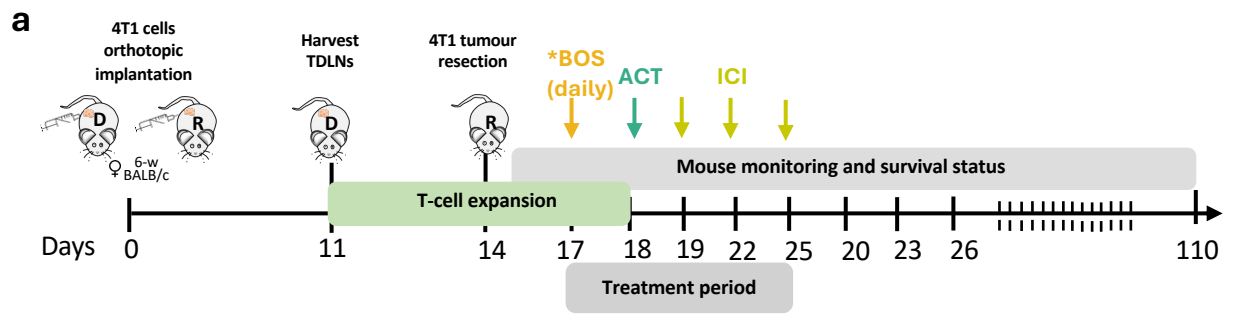

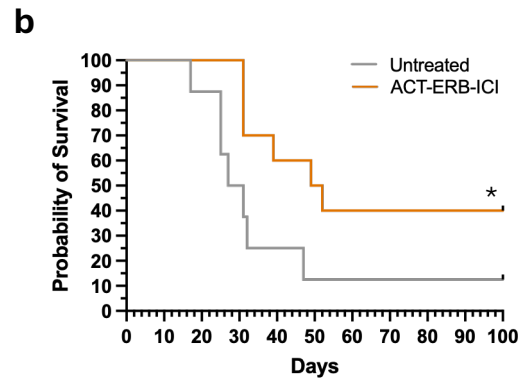

**Fig. S7: Post-operative ACT, endothelin receptor blockade and immune checkpoint inhibition prolong survival following surgical resection of established 4T1 tumours.**

**(a)** Experimental schematic. BALB/c mice bearing established 4T1 tumours underwent surgical resection of the primary tumour on Day 14 after tumour inoculation. Following surgery, mice received either no further treatment (control) or ACT combined with endothelin receptor blockade (ERB) and immune checkpoint inhibition (ICI). Survival was monitored until humane endpoint or study termination.

**(b)** Kaplan-Meier survival analysis of mice receiving surgery alone or post-operative ACT-ERB-ICI therapy. Post-operative ACT-ERB-ICI significantly prolonged survival compared with surgery alone (log-rank Mantel-Cox test, $P = 0.0398$). Four of ten treated mice remained alive at Day 100 compared with one of eight evaluable control animals. Two mice initially assigned to the control group were excluded from survival analysis because of post-operative wound-healing complications unrelated to tumour progression.
